# Synthesis of water-soluble and gastrointestinal transit-resistant FA-FOS conjugate for targeted delivery to the colon: pharmacokinetics, pharmacodynamics, efficacy

**DOI:** 10.1101/2023.05.05.539569

**Authors:** Eldin M Johnson, Joo-Won Suh

## Abstract

Ferulic acid is known to be a water-insoluble compound present in many fruits and vegetables and is known to possess antioxidant, anti-cancer, and anti-inflammatory properties. They are quickly absorbed in the stomach and metabolized in the liver. Their colonic exposure is found to be low due to their quick absorption and metabolism in the upper gastrointestinal tract, and due to this reason, only a small fraction of FA found in a bound form is associated with the insoluble and soluble fiber of the food matrix reaching the colon. Here we describe the synthesis and characterization of ferulic acid (FA) bound to fructo oligosaccharide (FOS) rendering the resultant FA-FOS conjugate water soluble, resistant to gastrointestinal digestion and absorption, along with the capability to deliver a therapeutically meaningful dose of FA to the large intestine. Free FA is released from FA-FOS conjugate by the digestive action of gut microflora, and the pharmacokinetic profile and pharmacodynamics are evaluated in a rat model. The efficacy of FA-FOS conjugate in the delivery of FA to the large intestine and its accumulation in tumours were evaluated in colitis induced colon cancer model and their efficacy through plasma bioavailability is determined in xenograft mice model carrying tumour from human colon cancer cells. The accumulation of FA derived from FA-FOS conjugate in the tumour was demonstrated by the MALDI imaging technique. The major metabolites of FA-FOS conjugate in plasma were determined through a data-dependent MS/MS experiment of precursor ion scan, utilizing triple quad (QTRAP) equipped LC-MS.

## 1 Introduction

Ferulic acid (4-hydroxy-3-methoxycinnamic acid, FA) is a phenolic acid that is found in abundance throughout the plant kingdom like in vegetables, cereals, and fruits. FA is known to possess potent antioxidant activity (1,2), They are found to be effective against cancer both in-vitro (3,4) and in-vivo (5–7) and are found to be completely safe even without any toxic side effects even at higher concentration, the LD_50_ for the male rat is 2.44 g/Kg while that of mice was 3.2 g/kg (8,9). They are also found to be effective as an adjuvant agent along with chemotherapy and radiotherapy (10). They are known to stimulate the immune system (11)and are found to have anti-inflammatory effect (12). Ferulic acid is reported to have anti-cancer activity on colo-rectal cancer (13–15). The anti-colo-rectal cancer activity of FA is due to its ability to regulate cell growth and proliferation, scavenging of free radicals, stimulatory effect on cytoprotective enzymes, and inhibition of cytotoxic reaction (8). Oral administration of FA is known to have poor gastrointestinal stability and low pharmacokinetic profile owing to its short plasma half-life and poor bioavailability limiting its clinical usage. On oral administration of FA, more than 75% of FA is absorbed by the gastrointestinal (GI) tract within 25 min and rapidly eliminated from the system as conjugated forms through bile and urine (16,17). On initial absorption into the gastrointestinal tract, FA is quickly metabolized in the liver and less than 9- 20% of total FA administered was available as free FA in the plasma, bile, and urine (16–20). Unmodified free FA found in urine accounts for only 4-5% of ingested FA both in rodents and in humans (18,20), the urinary excretion happens within 1.5 h after administration in rats (20) and in humans, the excretion of FA happens slowly with the plateau between 7-9 h after oral administration (18). Also, i.v (intra venous) administration of FA resulted in much higher levels of FA in the urine suggesting rapid filtration and excretion of free FA by the kidneys denoting low systemic exposure of FA. Serum albumin in the plasma is the major carrier of FA and it is estimated that free FA has 70% plasma protein binding as detected in human serum albumin (21,22). While only 4% of administered FA was observed in gastric mucosa (17), it was also found that almost 49% of perfused FA was distributed in the liver and peripheral tissues (23) and the concentration of the free FA decreased in most tissues just within 60 min of administration (24).

The rapid elimination of FA is influenced by the conjugation of FA in the liver, it was found that the bound form of FA (bound to arabinoxylans and xylans of the plant cell wall) is more bioavailable in the large intestine and the elimination rate from plasma was almost 15-fold slower than that of pure molecule (19,20). Previous studies also showed that the release of free FA from bound form happens only in the colon and is more bioavailable in the large intestine than administering free FA (19) and the released free FA was efficiently transported through the colonic epithelium without much metabolic transformation into feruloyl-glucuronide or sulphate (25). Among the cereal grains, refined corn bran has the highest FA content which is approximately 2.6-3.3 g/100 g of FA which has a serving size of 5 g, equivalent to 130-151 mg of FA per serving (26). It was estimated that the total amount of FA intake through the consumption of cereals, fruits, vegetables, coffee, etc., could add up to 150-250 mg/day (21). The FA that was found to be bound to arabinose or arabinoxylan in corn bran showed poor bioavailability than free FA(19). FA from corn bran has lower absorbability since they have FA bound to more complex heteroxylan (27).

In the treatment of colonic ailments, the drug must reach the colon without being absorbed by the small intestine and evade the first-pass metabolism by the liver and enzymatic action of the upper GI tract for the sufficient quantity (therapeutically relevant quantity) of the drug to reach the colon. Also, the presence of a meagre amount of feruloylated oligo saccharide or complex polysaccharide in the natural source like wheat/corn bran (1-3% bound ferulate)(28) results in a practically impossible volume of meal to be ingested to achieve a therapeutically relevant dose of FA in the large intestine and maintain the systemic exposure of FA for a reasonable time period. Also, the extraction and purification of such compounds from natural sources are highly uneconomical and non-feasible processes on a larger scale. Thus, from the prior research evidence, the present research was aimed at developing a conjugate between ferulic acid and fructo oligosaccharides (with varying degree of polymerization from n= 4-10) to deliver the active molecule FA directly into the large intestine by-passing the first pass metabolism and thereby achieving good bioavailability of free FA in the colon and maintaining the therapeutic relevant concentration of FA throughout the large intestine during the treatment intervention.

## 2 Materials and methods

### 2.1 Characterization

#### 2.1.1 Conjugation of ferulic acid onto fructo oligosaccharide and purification

All the chemicals and solvents were purchased from Sigma Aldrich unless otherwise mentioned. The fructo oligosaccharide was sourced from chicory which is composed of more than 90% fructo oligosaccharide and inulin, with an average degree of polymerization of > 9. The single pot two-step grafting of ferulic acid onto fructo oligosaccharide reaction condition and downstream processing were the same as that of our previous published work (29) except for some modification in the molar ratio of starting materials and esterifying agent which is maintained at 04: 04: 01 for trans-Ferulic acid: Carbonyl di-imidazole: Fructo oligosaccharide for the synthesis of water-soluble FA FOS II. After the completion of the reaction, the mixture was cooled to room temperature and the ferulic acid fructo oligosaccharide conjugate (FA FOS II) was precipitated by using three volumes of ice-cold isopropanol, the precipitate was washed several times with ethanol to remove the unreacted ferulic acid, the FA FOS II conjugates were dissolved in several washes of 50% ethanol and pooled. The pooled FA FOS II extract is centrifuged to remove any un-dissolved impurities carried over. The pooled FA FOS II washings were reduced to 35% of volume, in a rotary vacuum evaporator at 40°C. The resultant mixture is lyophilized to get a homogenous powder of pure ferulic acid oligosaccharide conjugate (FA FOS II). The purity of FA FOS II concerning unreacted FA and FOS is ensured by using HPLC equipped with a refractive index detector and TLC analysis.

#### 2.1.2 Degree of substitution with Ferulic acid

The degree of substitution (DS) is the average number of substituent groups (Ferulic acid) attached per base unit (fructo oligosaccharide). This DS value is experimentally determined by weighing a known amount of FA FOS II conjugates and adding 2M NaOH and incubating for two hours to release the ferulate group from the conjugate. The released ferulate group is then acidified to free ferulic acid by the addition of 6N HCl and adjusting the pH to 2.0. The free ferulic acid is then extracted from the aqueous solution by using three volumes of ethyl acetate three times. The pooled ethyl acetate extracts are then evaporated to dryness and re-suspended in a known volume of ethanol. The amount of the released ferulic acid is then quantified by using HPLC equipped with a diode array detector. The degree of substitution is then calculated by the formula:

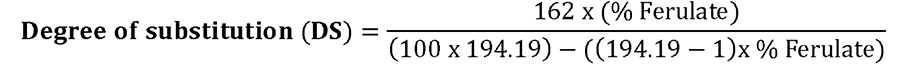

162 = Molecular weight of anhydrous glucose unit

194.19 = Molecular weight of Ferulic acid

#### 2.1.3 NMR analysis

The purified FA FOS II (35-40 mg) was dissolved in DMSO-d6 and scanned for overnight in 900 MHz Nuclear Magnetic Resonance spectrometer, AVANCE NEO 900 (Bruker) at Korea Basic Science Institute (KBSI), Ochang Center, Korea. The 1H1, 13C, 13C-DEPT −135° (Distortionless enhancement by polarization transfer), HSQC, and HMBC experiments were performed for elucidating the structure of the synthesized molecule.

#### 2.1.4 ESI MS

The ESI MS spectra of FA FOS II were collected using Triple Quadrupole LC-Mass spectrometry (Finnigan TSQ Quantum Ultra EMR) at KBSI, Seoul center. The sample was filtered using a 0.22 µm membrane filter and directly infused into the mass analyzer using an automated syringe at a flow rate of 20µL/min after dissolving 1 mg/mL of FA FOS II in 0.1% formic acid. The spectra were collected in negative ion mode with a spray voltage of 3000 mV, sheath gas pressure of 10 psi, auxiliary gas pressure of 5 psi, and a capillary temperature of 300° C.

#### 2.1.5 Fourier transform infrared spectroscopy (FTIR) analysis

FTIR analysis was performed for FA, FOS, and FA FOS II to validate the grafting of FA onto FOS. The powder samples were mixed with dry potassium bromide (KBr) in the ratio of 1:200 (wt/wt) and pelleted by using a hydraulic press. The FTIR spectra of the prepared discs were recorded using an IR spectrometer (Varian 2000 FT-IR, Scimitar Series). The background spectrum was collected with clean KBr discs and this background noise was subtracted from the sample signal.

#### 2.1.6 X-ray diffraction studies

The X-ray diffraction pattern was analyzed by using Malvern Panalytical, AERIES 600 X-ray diffractometer with Cu as anode material and radiation K-α2/ K-α1 ratio of 0.5000 at 40 kV and 15 mA. The diffractometer was equipped with an automatic divergence slit and the scattered radiation was detected in an angular range of 5-50 (2θ), with a scanning speed of 2° (2θ)/min and step size of 0.022° (2θ).

#### 2.1.7 Zeta potential and particle size analysis

The Z-average diameter and ζ potential of FA FOS II were performed in neutral pH by dissolving in deionized water and diluted to a concentration of 0.5 mg/mL and determined by using dynamic light scattering (DLS). The measurements were performed at a fixed scattering angle of 173° using a Zetasizer Nano-ZS (Malvern, UK) at 25°C equipped with a He-Ne laser (λ = 633 nm). The average of triplicate measurements was calculated for each parameter.

#### 2.1.8 Thermogravimetric analysis

The thermogravimetric analysis (TGA) was performed in Mettler Toledo DSC 3 (TGA 2 star system). Samples from 5-10 mg were heated at a rate of 20° C/min from ambient temperature to 800° C. The purge gas used was nitrogen at a flow rate of 50 ml/min. The thermal stability was determined from the Tx% value which is the temperature corresponding to x% mass loss.

### 2.2 In-vitro stability and release

#### 2.2.1 Solubility and stability testing

The solubility was tested by suspending 10 mg FA FOS II in different solvents and their mixtures with varying polarity and in ionic liquids. The suspension was mixed and the solubilization was attempted using an ultrasonic water bath set at 60° C. The stability of the FA FOS II conjugate in simulated salivary and gastrointestinal fluid at the different time points of incubation at 37° C from 0 min, 15 min, 30 min, 1 hr up to 4 hr were investigated. The composition of simulated fluids is as described by Bove et al (30) with modifications as described below. The composition of the simulated salivary fluid is CaCl_2_.2H_2_O at 0.0228%, NaCl at 0.1017%, Na_2_HPO_4_ at 0.0204%, MgCl_2_ at 0.0061%, K_2_CO_3_ at 0.0603%, Na_2_HPO_4_ at 0.0273%, α-amylase at 2 units/ml, lysozyme at 0.0015% and pH is adjusted to 7.4. The simulated fasted state gastric fluid is composed of sodium taurocholate 0.0043%, lecithin 0.0051%, pepsin 0.01 %, lysozyme 0.01 %, and NaCl 0.09% adjusted to the pH of 1.3 using HCl. Simulated fasted state intestinal fluid is composed of sodium taurocholate 0.0118%, lecithin 0.0465%, Malic acid 0.2219%, NaCl 0.2%, Pancreatin (4 USP/mg) 0.050% adjusted to pH=6.5. The degradation or digestion is monitored by checking for the presence of the ferulate group released into the simulated fluids during the incubation period. The extraction of the ferulate group from the simulated fluid involves the acidification of the simulated fluid containing FA FOS conjugates and without FA FOS conjugates (negative control) to pH= 2.0 using 6N HCl and the addition of three volumes of ethyl acetate and extraction was carried out three times as mentioned above. The pooled extracts were evaporated to dryness and dissolved in ethanol and the presence of any ferulate group released during the digestion was determined by using HPLC equipped with a diode array detector.

#### 2.2.2 Stability under the action of enteric enzymes and gut microbiota

About 250 mg of FA FOS II conjugate was added with 50 mg of Driselase and 50 mg of Protease M Amano from Amano enzymes, Japan along with 50 µL of DEPOL 670L and DEPOL 740L each from Biocatalyst, UK along with 0.02% Sodium Azide were dissolved in 5 ml MOPS buffer of pH= 6.0 and incubated at constant agitation for 12, 24, 36, 48 h at 37° C. The above enzyme mix is a cocktail of cellulase, endo-galactouranase, endo-carbohydrase, exo-glucosidase, and feruloyl esterase which are the enzymes produced by the enteric bacteria living in the colon. After the enzymatic treatment process, the released free ferulic acid from FA FOS conjugates was quantified by acidifying the enzymatic mixture to pH=2.0 using 6N HCl and extracting with three volumes of ethyl acetate, pooled and analyzed with HPLC.

The FA FOS conjugates were digested under simulated colonic conditions by using human microbiota as inoculum. Briefly sterilized anaerobic minimal basal medium of 10 ml according to (31) was added with 0.05g of FA FOS conjugates in a head space vial and added with 1% human feces w/v under anaerobic condition in an Anaerobic chamber and sealed under anaerobic condition. The vials were incubated at 37° C for 12, 24, 36, and 48 hours with intermittent agitation. After the termination of incubation, the fermentation medium is centrifuged at 10000x G for 10 min at 4° C and the supernatant was collected. The extraction and quantification of free ferulate released from FA FOS conjugate were performed as described above.

#### 2.2.3 LC MS/MS analysis and identification of the metabolite of FA FOS II

The HPLC analysis was carried out by using a C_18_ Nucleosil Column with a flow rate of 1 ml/min at a column temperature of 25° C and the detection was carried out at 280nm using a diode array detector. The solvent system used is mobile phase A which is composed of Acetonitrile: H_2_O: Formic acid at a ratio of 1.5: 7: 0.0312 (v/v) at pH = 2.33 and mobile phase B which is composed of Acetonitrile: H_2_O: Formic acid at a ratio of 7: 1.5: 0.0312 (v/v) at pH= 2.66. The gradient program was as follows: Initial 0-10 min are mobile phase A, the percentage of mobile phase B is increased to 100% over a gradient from 10-30 min followed by a holding time of 10 min with 100% mobile phase B and later the column is regenerated back with mobile phase A 100% over a gradient time of 10 min.

The Q-TOF MS operating conditions were Electrospray ionization (ESI) in negative mode, desolvation temperature was 300° C, source temperature 120° C. Capillary voltage = 1.2 kV, Sample cone voltage = 45 V, Extraction cone voltage = 4 V, Cone gas flow = 50 L/hr, desolvation gas flow 900 L/hr, Collision energy for MS/MS set to 20 V.

### 2.3 In-vivo studies in an animal model

#### 2.3.1 Pharmacokinetic and pharmacodynamics study in Rat

The pharmacokinetic (PK) study was carried out in Sprague-Dawley rats weighing 180-250g (7-9 weeks old) provided by Charles River Laboratories. Space allocation for the individual animal was 47 x 25 x 21 cm. All animals were maintained in a controlled temperature (20 - 24°C) and humidity (30% - 70%) environment with 12 hours of light/dark cycles. Free access to a standard lab diet (Oriental Yeast Co., Ltd., Japan) and autoclaved tap water was granted. All aspects of this work including housing, experimentation, and animal disposal were performed following the “Guide for the Care and Use of Laboratory Animals: Eighth Edition*”* (National Academies Press, Washington, D.C., 2011) in an animal facility in Taiwan. In addition, the animal care and use protocol was reviewed and approved by the IACUC at Pharmacology Discovery Services Taiwan, Ltd. The experimental groups were:

i. Oral (P.O) administration of FA 100 mg/kg body weight in 4% DMSO added with 0.5% tween 80, as a suspension with a dosing volume of 10 ml/kg for PO,
ii. Oral (P.O) administration of FA FOS II 3.89 g/kg (100 mg/kg free FA equivalent) dissolved in water,
iii. Intravenous administration (IV) of FA 100 mg/kg, in a formulation of 8% DMSO added with 5% Solutol^®^ HS-15 in Phosphate buffered saline (PBS) as suspension, with a dosing volume of 5 ml/kg.

The formulations were prepared fresh before administration. Each dosage group had three replicate animals and one animal each dosed with PBS and water respectively served as blank for IV and PO. The blood samples were collected at 0, 0.083, 0.167, 0.25, 0.5, 1, 3, 6, 8, 12, and 24 h post dosage. A few days before the compound administration, animals were initially anesthetized with pentobarbital [50 mg/kg, IP] injection for jugular catheter implantation. A single dose of meloxicam [1 mg/kg, subcutaneously (SC)] was given as an analgesic to relieve the postoperative pain. Blood aliquots (300- 400 μL) were collected from jugular vein catheterized rats into tubes coated with lithium heparin at the specified time intervals. The tubes were mixed gently and kept on ice and then centrifuged at 2,500 ×g for 15 minutes at 4°C, within 1 hour after collection.

The plasma and the dosing solution samples were analyzed by LC-MS/MS. The detailed chromatographic conditions, bioanalytical methods, and acceptance criteria are summarized as follows: LC MS/MS instrument used was SCIEX Triple Quad^TM^ 5500+, electrospray ionization in positive ion mode, multiple reaction monitoring (MRM) scanning method is used, MRM of analyte is 193/133.1 (Q1/Q3) and the internal standard used was Oxybutynin with MRM 358.3/142.32, the analyte FA is quantified using peak area ratio of the known standard and weighted linear regression of the same. The column used was Agilent Poroshell 120 EC-C18 column of 3.0×50 mm and a pore size of 2.7 µm at a column temperature of 40° C, the injection volume is 5 µL and run time is 3 min using 0.2% v/v formic acid in water and 0.2% v/v formic acid in acetonitrile. The initial acetonitrile concentration of 85% is reduced to 30% over a gradient time of 1 min followed by a hold for 1 min and increased back to 85% over a gradient time of 1 min. About 20 µL of rat plasma is mixed with 300 µL of methanol containing 0.01 ng/µL internal standard and vortexed for 1 min followed by centrifugation at 3000xG for 5 min. The supernatant of 200 µL was added with 300 µL 0.2% formic acid in water and injected into LC MS/MS analyzer.

For determining the various metabolites of FA in the plasma, the precursor ion of the parent is fixed in Q3 and the parent ion is scanned in Q1 in this data-dependent experiment. All the metabolites of FA are known to contain the fragment ion/ precursor ion 133.9 and hence the Q3 of the triple quad detector is fixed for the selection of fragment ion 133.9 and whenever the selected Q3 is detected, the instrument is triggered to scan the Q1 to screen for the parent molecule giving rise to the product/precursor fragment detected in Q3 and full MS and MS/MS data is collected and the metabolite is identified. The abundance of the molecule/metabolite of FA in the plasma is derived from the XIC (Extracted ion chromatogram) of the screened molecule’s precursor ion, intensity in the plasma.

#### 2.3.2 Efficacy studies in xenograft mice model

The anti-tumor efficacy of FA FOS II through its bioavailability and its metabolites in the blood plasma was investigated by using Xenograft mice carrying HT-29 tumour. The cell line was cultivated in RPMI 1640 medium with 10% FBS under 5% CO2 atmosphere and the cells were harvested and diluted to 1×10^7^ cells/0.2 ml/mice and injected into the right flank subcutaneous region of Specific Pathogen Free (SPF) BALB/c Nude mice (CAnN.Cg-Foxn1^nu^/CrljOri) from OrientBio Inc., Korea and grown until the tumour volume reached 150-200 mm^3^. Animals were provided with Teklad Certified irradiated global 18% protein rodent diet (Envigo, UK) and sterile water with free access to them. The wood chip bedding was autoclaved before use.

The BALB/c Nude mice carrying HT-29 tumour were randomly allotted to two treatment groups + 1 Naïve group:

i. Vehicle Control group-no treatment,
ii. FA FOS II treatment group – 3.890 mg/kg body weight, vehicle-distilled water, P.O administration not more than 10ml/kg, administered every day for four weeks
iii. 5-Fluorouracil (5-FU) treatment group-Initial dose = 30 mg/kg body weight, vehicle- 0.9% saline, administered I.P (Intra peritoneal), not more than 0.2ml once a day for 5 consecutive days a, followed by rest for 1 week and I.P at a dose of 15 mg/kg twice a week for the next two weeks. This treatment regime was chosen because of the adverse side effects of 5-FU observed during the study.
iv. Naïve group-without tumour and treatment

The clinical signs, body weight, and tumour volume are observed twice a week for four weeks during the treatment period. After the elapse of the dosing of the animals for four weeks the animals were anesthetized with isoflurane and the blood samples were collected from the cephalic vein into heparin tubes, immediately plasma was harvested and stored at −80° C. The animals were sacrificed by exsanguination from the posterior vena cava and aorta. Tumours were then removed, weighed, and sent for immunohistochemistry (IHC) analysis after fixing in 10% neutral buffered formalin solution for 24h and embedded in paraffin blocks using an automated tissue processor (Shandon Citadel 2000, Thermo Scientific, USA) and embedding unit (Shandon Histostar, Thermo Scientific, USA). About 3-4 µm sections were prepared using an automated microtome (RM2255, Leica Biosystems, Germany) and the sections were stained with hematoxylin and eosin (H&E) staining, immunohistochemistry, and immunofluorescence staining accordingly using BenchMark ultra, Ventana IHC/ISH auto-Stainer system of Roche diagnostics, Switzerland in a well standardized automated condition so that all the staining, counter staining and washing steps are all homogenous for all the samples. The slide was scanned using Pannoramic 250 Flash III, 3DHISTECH Ltd., Budapest, Hungary in a multi-focus mode at 200x and analyzed using QuPath 0.3.2 software to determine the Dab staining, fluorescence intensity, and other histopathological parameters. The study was approved by the Institutional Animal Care and Use Committee (IACUC) of Chemon Inc., Gyeonggi Bio Center (Serial# 2019-08-002).

#### 2.3.3 Efficacy studies in colitis-induced colon cancer mice model

The strain of mice used in the induced colitis-associated colon cancer mice model is C57BL/6, 5-6 weeks-old female mice were randomly assigned to four groups of seven mice each. Initially, the mice were injected with a single intraperitoneal injection of azoxymethane (AOM) 8 mg/Kg bodyweight of mice followed by administration of AOM after dissolving in water via oral gavage per oral (P.O) once per day for a week. Following AOM administration the mice were fed with 2% DSS for a period of one week, dissolved in drinking water. After a brief resting period of one week, the mice were fed with 1.5% DSS in drinking water for another week followed by a week of rest and subsequent administration of 1% DSS for another week. This is followed by a week’s rest and the FA FOS conjugates were administered at the indicated dosage below for four weeks.

The experimental groups were:

i. Naïve group (Mice without colon cancer induction)
ii. Control group (AOM-DSS induced tumour bearing mice),
iii. FA treatment group – 100 mg/Kg, P.O, vehicle-Water
iv. FA FOS II treatment group – 3890 mg/kg (equivalent to 100 mg/Kg free FA), vehicle-water

After the termination of 4 weeks, the mice were anesthetized under isoflurane and the blood was extracted through the cardiac puncture into heparin-coated tubes, the mice were euthanized by cervical dislocation, and the organs like the kidney, liver, spleen, and thymus, colon and femoral bone were extracted and used for further analysis. This experimental procedure is done according to the regulation and approval of the Institutional Animal Care and Use Committee (IACUC) of Gachon Medical Research Institute, Le Gil Ya cancer research center, Incheon, Korea.

#### 2.3.4 MALDI imaging and confirmation of targeted delivery

The full-length swiss rolled colon from Colitis induced colon cancer model, the xenograft model’s excised tumour, liver, and kidney were snap frozen in liquid nitrogen and stored at −80° C. They were freshly processed for MADLI imaging analysis and to determine the localization and delivery of the FA/ FA metabolite at the target site by FA FOS II. The tissue samples snap frozen in liquid nitrogen were embedded in carboxymethyl cellulose and sectioned in cryostat set at −20° C and tissue sections of 10 µm were collected on a conductive ITO glass slide briefly warmed using a finger at the non-conductive side and CMC is removed. The dihydroxybenzoic acid (DHB) matrix was used for MALDI analysis in Positive ion mode using Fourier transform ion cyclotron resonance (FT-ICR) and full spectra were collected at a mass range of 104 m/z to 1000 m/z.

## 3 Results and discussion

### 3.1 Characterization of FA FOS II

#### 3.1.1 Degree of substitution

The degree of substitution with FA was determined as previously described in (29). The percentage composition of FA in FA FOS II was 2.575 % wt/wt and the degree of substitution was found to be 0.02. They are readily soluble in water, DMSO and as aqueous mixtures in organic solvents like acetonitrile.

#### 3.1.2 NMR analysis

The structural features of the FA FOS II were elucidated by using NMR analysis. In the ^1^H NMR, the characteristic peaks of the ferulate group were observed between 6.5 to 7.8 ppm and the sugar moiety was observed between 2 to 5 ppm. The intensity of the signals from ferulic acid relative to that of the carbohydrate region shows that the compound has mainly a single ferulic acid moiety attached to each molecule of FOS. The multiplets observed were 1H NMR (900 MHz, DMSO) δ 11.94 (s, 1H), 9.60 (s, 1H), 7.61 (s, 1H), 6.98 (s, 1H), 6.75 (t, *J* = 6.7 Hz, 1H), 5.14 (s, 9H), 4.70 (s, 9H), 4.63 (s, 1H), 4.58 (s, 9H), 4.36 (s, 1H), 4.07 – 4.04 (m, 1H), 4.04 – 3.96 (m, 7H), 3.96 – 3.93 (m, 1H), 3.78 (dq, *J* = 16.3, 9.3, 8.2 Hz, 13H), 3.72 (s, 1H), 3.61 – 3.58 (m, 9H), 3.55 (d, *J* = 10.4 Hz, 9H), 3.52 – 3.49 (m, 2H), 3.48 – 3.45 (m, 17H), 3.45 – 3.41 (m, 7H), 3.35 (s, 4H), 3.34 (s, 1H), 2.46 (s, 1H). The protons from –OCH_3_ of ferulic acid were observed at 3.47 ppm, and the protons from –CHCHCOO of ferulic acid moiety involved in ester bond formation were observed at 6.75 ppm. The protons from the aromatic group of ferulic acid were observed between 6.98 to 7.3 ppm, and the weak peak at 9.60 ppm represents the phenolic OH. The sugar protons were observed between 3.43 to 5.14 ppm. 1H spectra with assignment in green can be found in Fig. 2a and the molecular structure along with numbering is represented in Fig. 1. The TOCSY spectra, HSQC & HMBC (1H-13C DEPT 135) along with their annotations were presented in the supplementary figure S1a, b, c, d, e. In the 13C NMR spectra the – OCH_3_ group was observed at 56.07 ppm the carbonyl group was observed at 166.65 ppm, and the aromatic and side chain of the feruloyl moiety was observed between 115 to 149 ppm. The carbons from the sugar moiety were observed between 61 to 103.3 ppm (Fig. 2b). The multiplets observed were 13C NMR (226 MHz, DMSO) δ 166.67, 149.37, 148.02, 147.96, 147.36, 145.82, 145.37, 135.24, 125.69, 125.59, 123.82, 123.49, 123.24, 118.00, 115.61, 115.55, 114.41, 114.11, 113.02, 111.27, 111.16, 104.02, 103.33, 103.32, 103.29, 103.21, 103.14, 92.17, 82.31, 82.07, 81.81, 81.65, 80.01, 78.77, 77.34, 77.08, 76.67, 76.51, 75.17, 74.70, 74.51, 74.41, 74.22, 72.99, 72.91, 71.76, 69.85, 65.54, 62.13, 61.73, 61.56, 61.13, 60.47, 56.07, 55.80, 55.75, 55.72, 40.43. The –CH_2_ from the sugar moiety was observed as inverted peaks between 51.9 and 61.9 ppm in the ^13^C DEPT 135 NMR spectra (Fig. 2c).

**Fig. 1.**
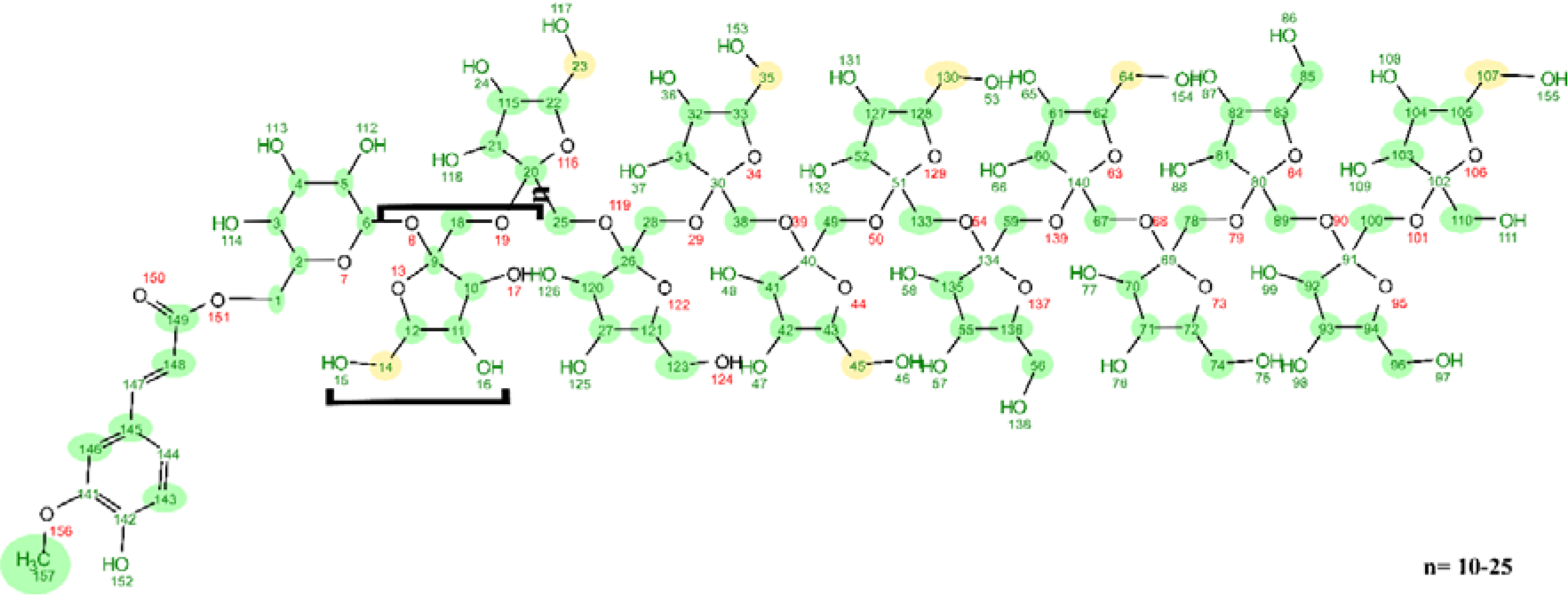
Elucidated structure of FA FOS II

**Fig. 2.**
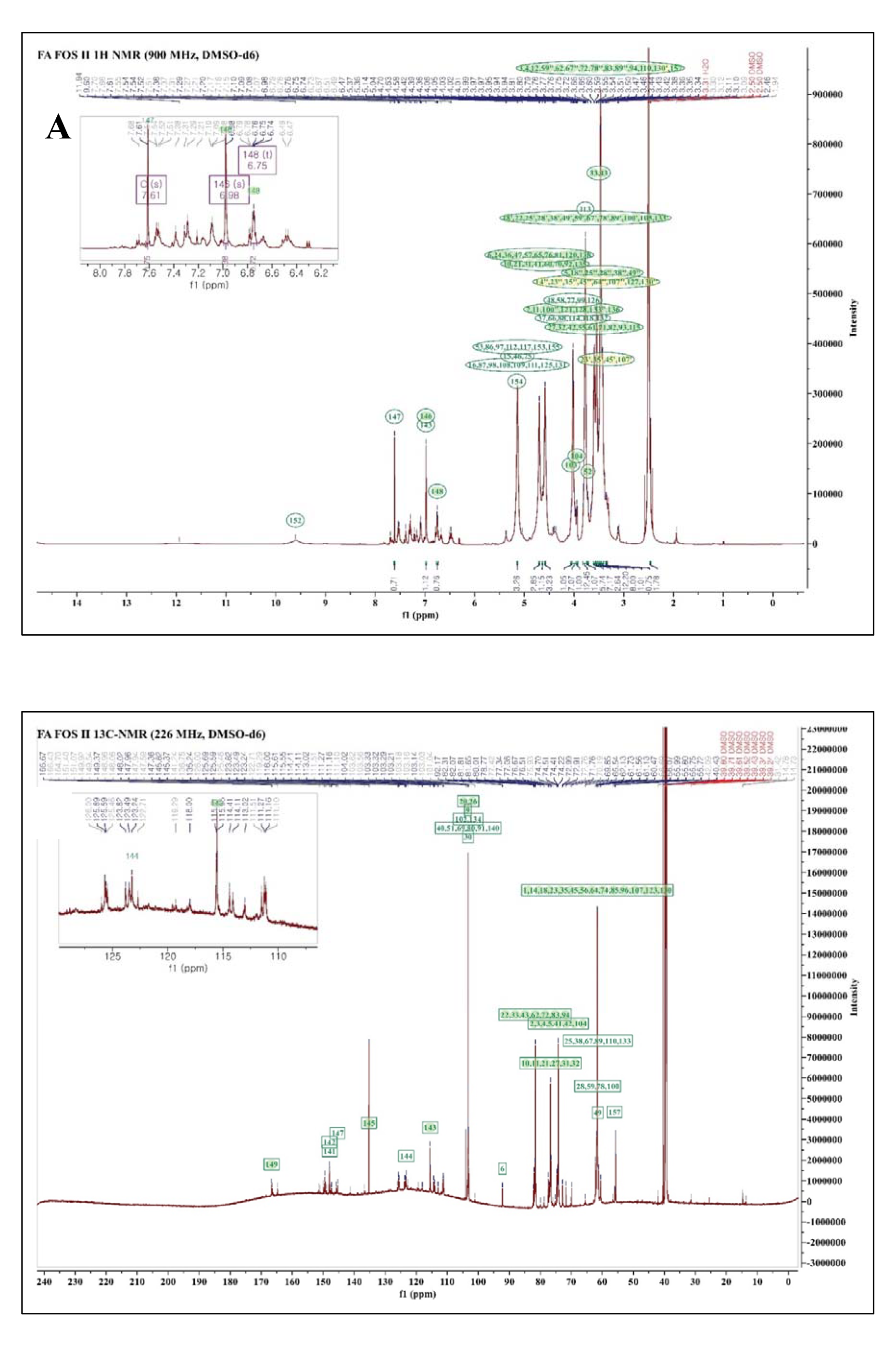
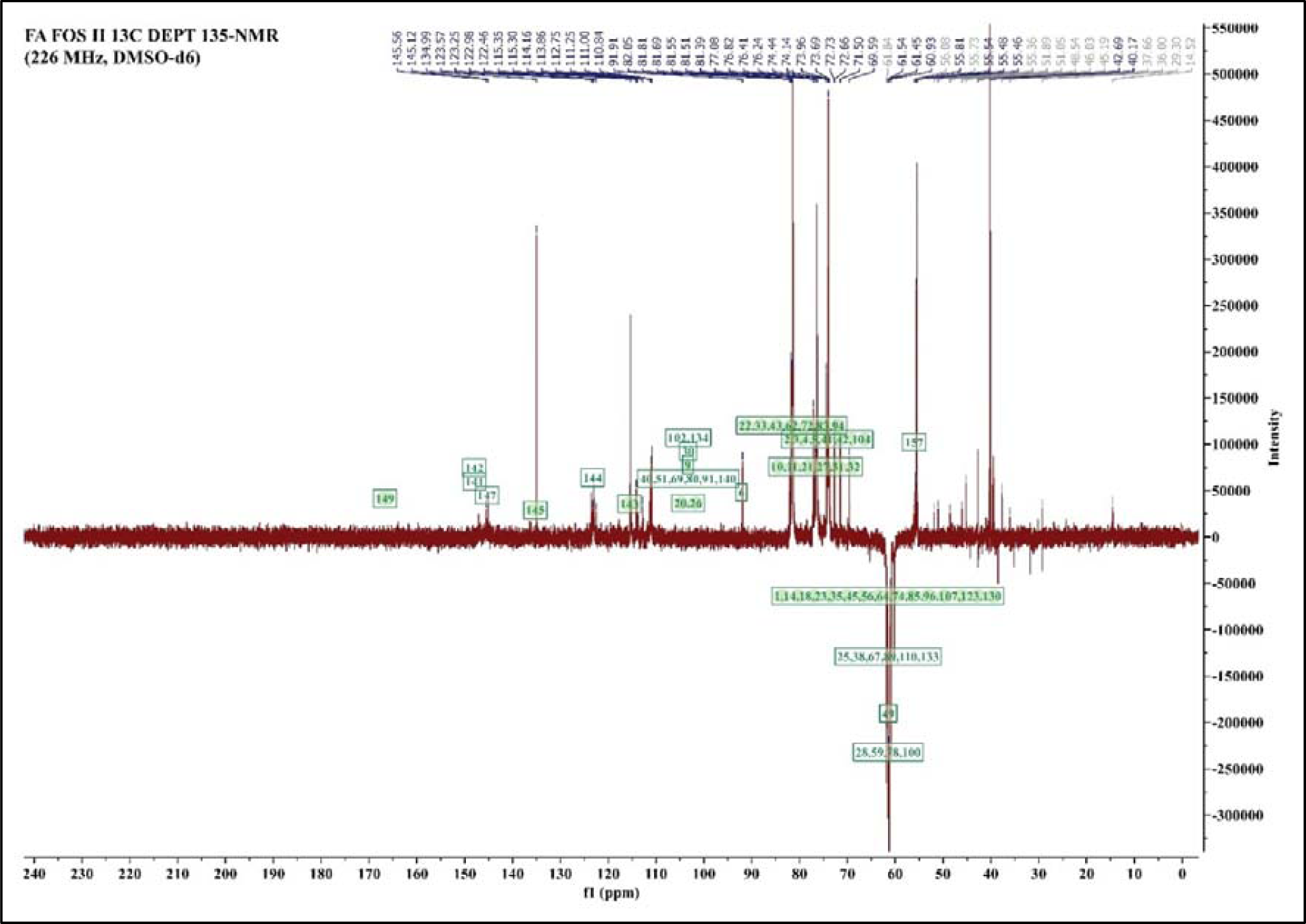
NMR analysis and annotation of FA FOS II (A) Proton of FA FOS II, inset picture-zoomed area from 6 to 8 ppm, (B) 13C NMR of FA FOS II, inset picture-zoomed shifts from 110 to 150 ppm, (C) 13C DEPT at 135° angle to differentiate between CH and CH_3_ which are found at a phase opposite to CH_2_

#### 3.1.3 ESI MS analysis

The ESI MS was performed to determine the degree of feruloylation and the position of the conjugation of ferulic acid onto the oligosaccharide molecule. ESI-MS using negative ion mode was performed, and full MS of the purified reaction product showed the presence of mono-feruloylated oligosaccharide, with the fructo oligosaccharide degree of polymerization ranging from n= 09-21. (Fig. 3)

**Fig. 3.**
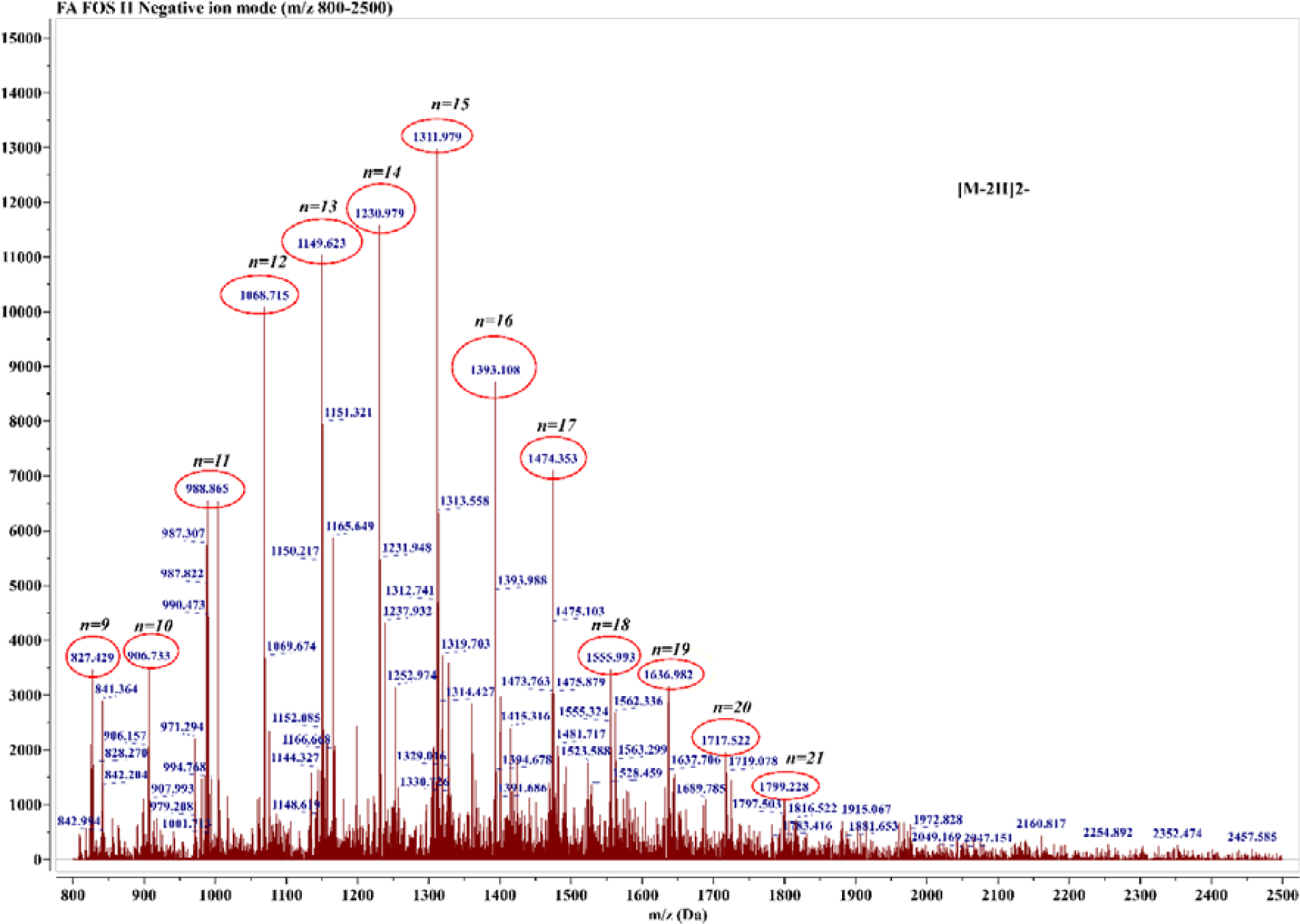
ESI MS of FA FOS II in negative ion mode, n represents the degree of polymerization of FOS moiety of the FA FOS II

#### 3.1.4 FTIR analysis

From the FTIR spectra (Fig. 4) the broad valley at 3400 cm^-1^ corresponding to the hydroxyl group in FOS is reduced in FA FOS II owing to the substitution of the OH group by involving in esterification with ferulic acid. The carbonyl signal at 1717 cm^-1^ in the FA FOS II indicates the successful esterification of FOS with the feruloyl group however this peak is smaller compared to that reported by (29) owing to the lesser degree of substitution with FA (32). The carboxylic acid C=O and O-H stretches corresponding to the peak at 1693 cm-1 and 2846 cm^-1^ respectively found in FA were absent in FA FOS II owing to esterification. The peaks 1625.6, 1593.8, 1515.7, and 1431.8 cm^-1^ corresponding to the C-C stretch characteristic of the aromatic ring are coming from FA substitution in the FA FOS II spectra.

**Fig. 4.**
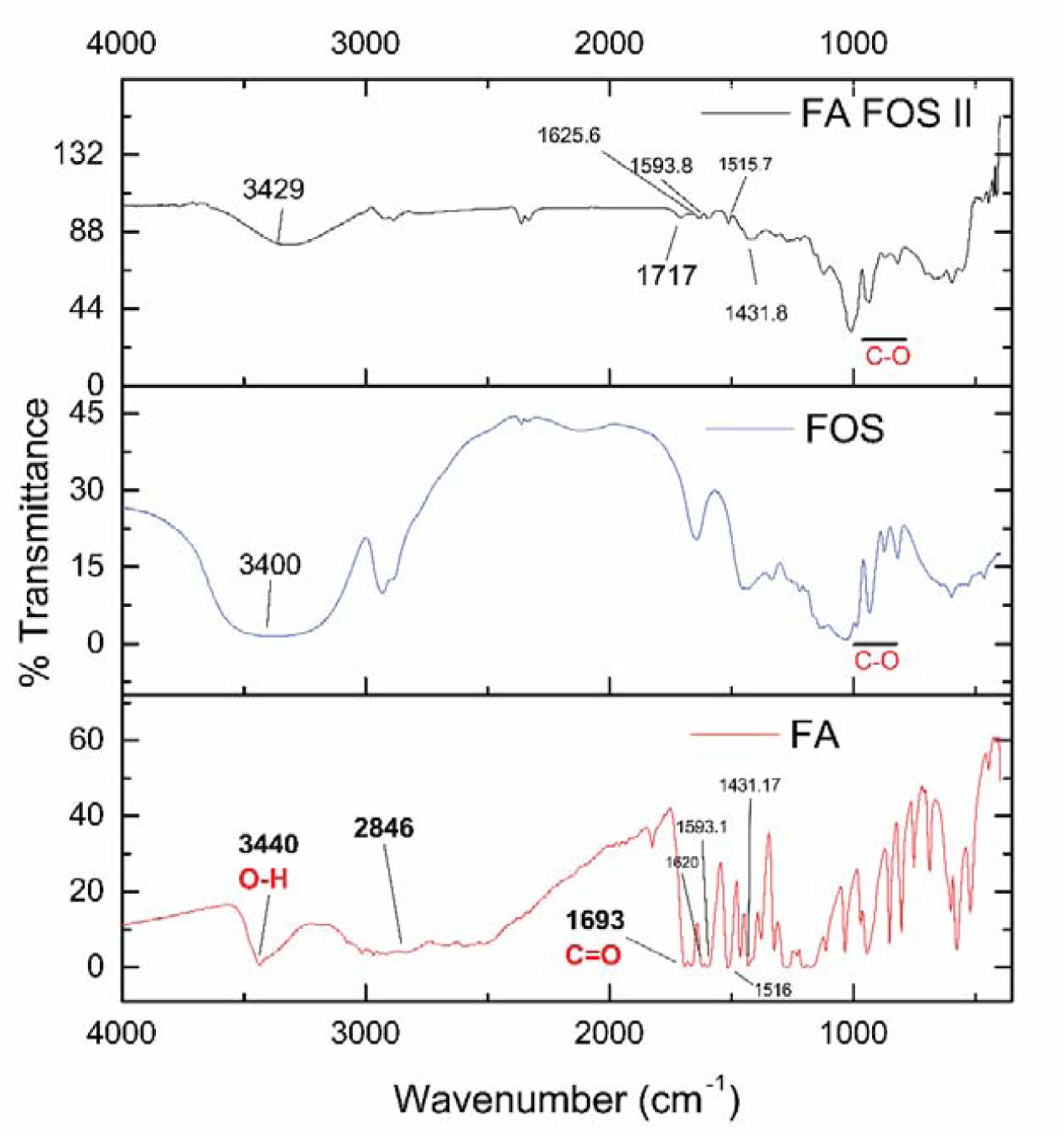
FTIR spectrum of FA FOS II, FOS, and FA

#### 3.1.5 X-ray Diffraction studies

The XRD pattern of FA FOS II conjugate shows amorphous or nano-crystalline nature with a weaker peak at 11.96° and a broad hump between 5°-45° with reduced intensity compared to that of FA and FOS (Fig. 5) suggesting nano-crystalline type material or smaller particle size, while the native FOS showed a semi-crystalline pattern with weaker peaks at 8°, 16.13°, 24.09°, 30.12°, 33.80° and stronger peaks at 12.07°, 17.65°, 21.72°. The FA FOS II did not show any crystalline peaks of FA which showed a highly ordered crystalline nature with sharp and strong peaks at 9.03°, 10.50°, 12.83°, 15.63°, 26.46°, 29.43° respectively. The results of XRD of FA FOS II suggest that the modification of native FOS with hydrophobic FA resulted in the disruption in the ordered structure of native FOS which could otherwise exclude water due to their three-dimensional ordered structure due to weak hydrogen bonds and water bridges (33,34). The hygroscopic nature of the newly synthesized FA FOS II can be explained by this phenomenon. However the higher degree of substitution of FOS with FA, they maintained the semi-crystalline nature as reported by us earlier (29), which may be due to the formation of disc-shaped spherulites/ aggregates in the aqueous media during the preparation process because of decrease in water solubility (35).

**Fig. 5.**
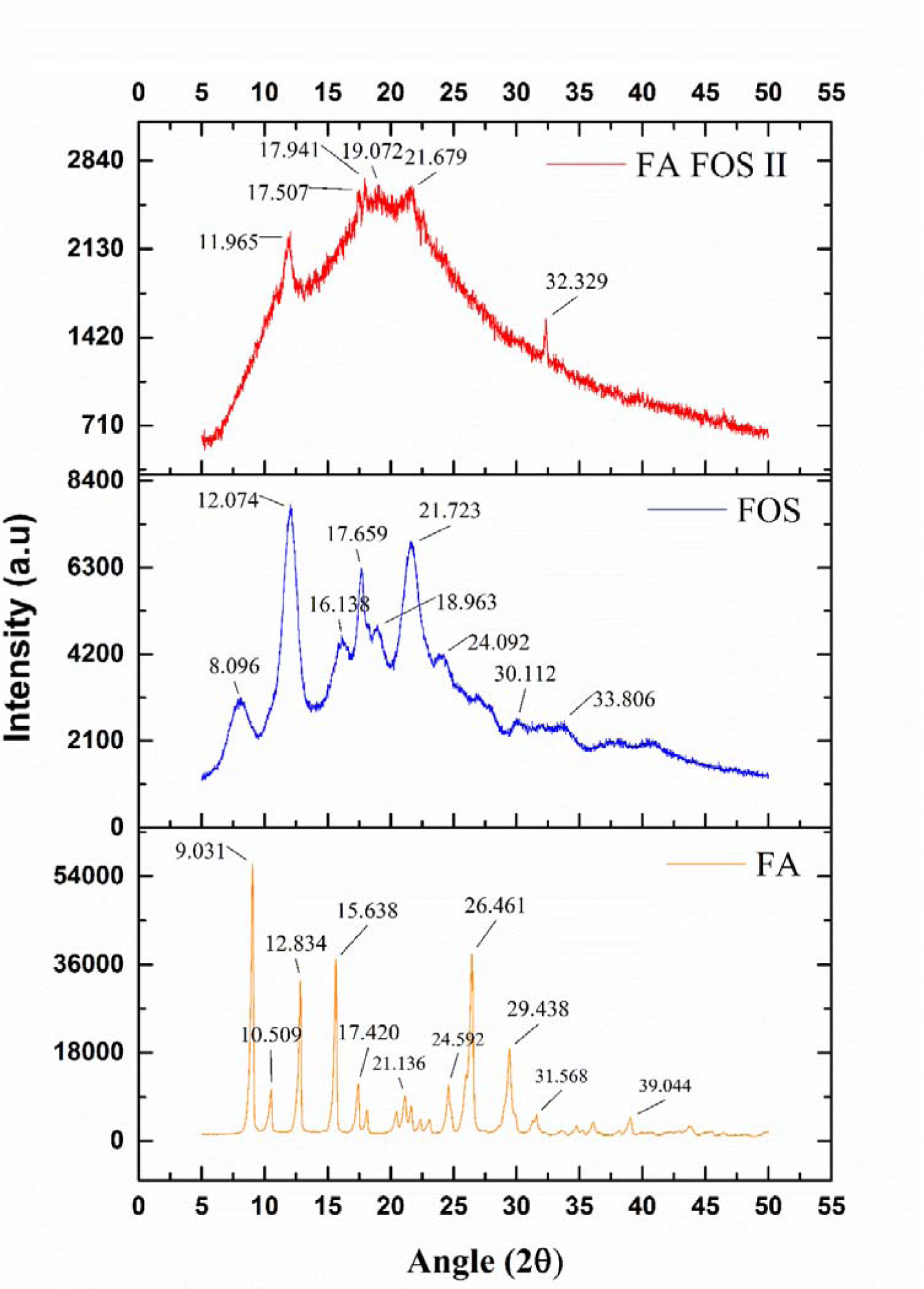
X-ray diffraction pattern of FA FOS II contrasted with that of FOS and FA

#### 3.1.6 Zeta potential and particle size analysis

The ζ-potential of FA FOS II dispersed in de-ionized water was found to be −12.1 ± 5.83 mV and conductivity of 0.0491 mS/cm. The average particle size (diameter) was found to be 204.6 ± 61.08 nm and a polydispersity index of 0.176 as measured by DLS (dynamic light scattering). The small particle size explains the nano-crystalline-like XRD pattern and reduced intensity of the peaks (Fig. 6). The FA FOS II seems to be forming ultrafine-grained structures when dispersed in water by hiding the hydrophobic FA (guest) well within the core isolating them from the hydrophilic outer shell (inclusion complex) which interacts with the water and thereby rendering the compound soluble (36).

**Fig. 6.**
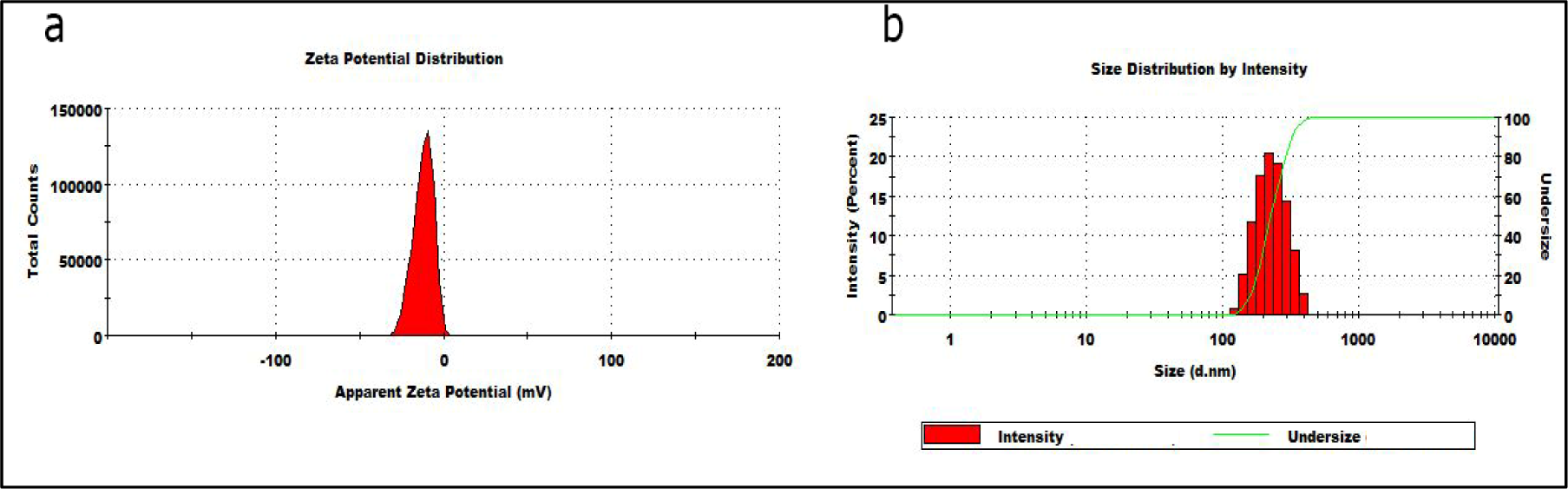
The ζ-potential (a) and particle size distribution (b) of FA FOS II

#### 3.1.7 Thermogravimetric analysis

The thermogravimetric analysis (TGA) was performed to determine the thermal stability of the compound and to identify the decomposition pattern. The TGA profile of FA FOS II shows three-stage weight loss at 141.11° C, 233.75° C, and 304.41° C. There was observed a 12.32% weight loss observed at the temperature range from 70° C to 140° C at the rate of loss of 0.004 mg/ml from 70° C - 100° C and at the rate of 0.03 mg/ml beyond 100° C, which might correspond to the water loss/moisture absorbed by the compound. There was a 43.62% reduction in weight at 233.75° C for FA FOS II due to the intermolecular and intra-molecular condensation of hydroxyls of FOS (29), beyond 240° C the loss in weight corresponds to the degradation of FA, this suggests that the compound might be resistant/durable on sterilization by autoclaving at 121° C. The synthesized compound FA FOS II was found to be more thermally stable than the native FOS and free FA but to a much lesser extent compared to the material with a high degree of substitution reported by us earlier (29).

**Fig. 7.**
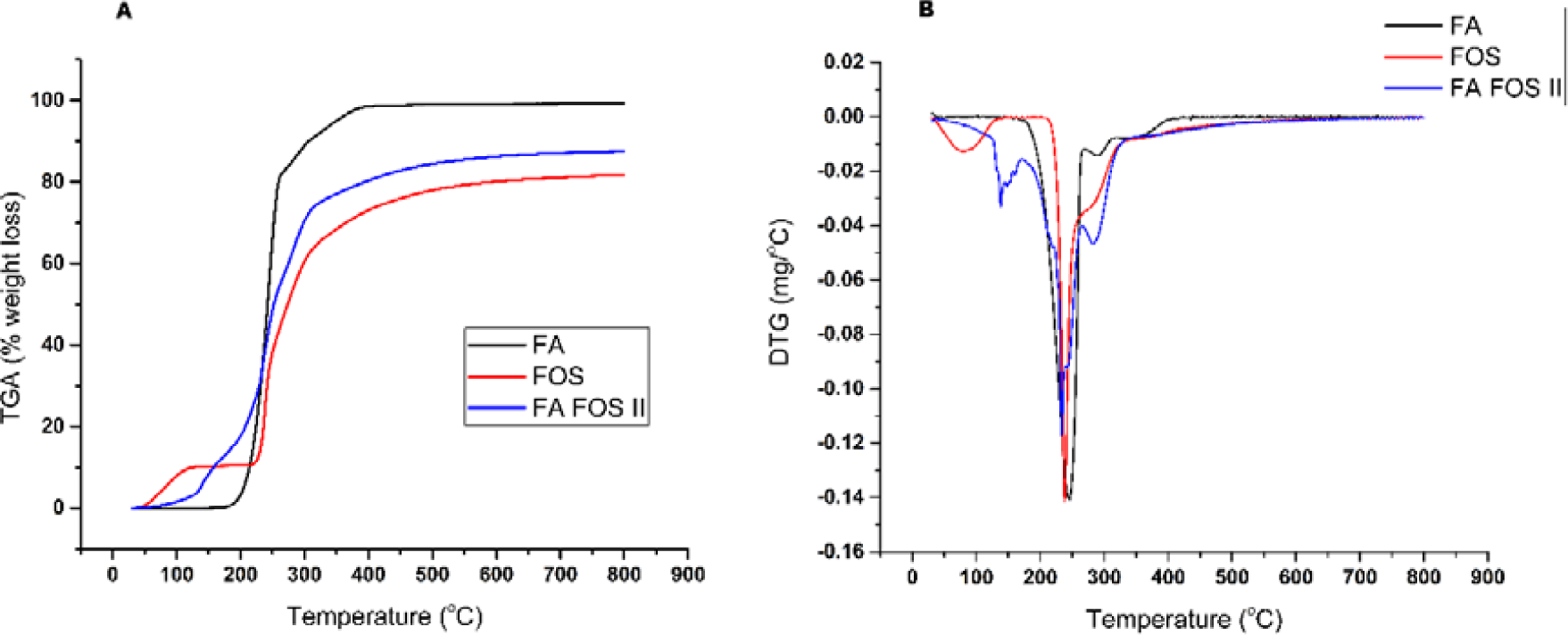
(A) TGA thermogram and (B) Derivative Thermogravimetric Curve of FA, FOS, and FA FOS II

### 3.2 Solubility and stability test of FA FOS conjugate

The FA-FOS II was found to be stable under simulated gastrointestinal fluids. But it was found that FA-FOS I is more resistant to the enteric enzyme and released the least free ferulic acid compared to that FA-FOS II which released about 45.61% of its FA content (29). The stability of the FA-FOS II was also tested under the simulated colonic condition with the human faecal sample as inoculum under anaerobic conditions. The FA FOS II was found to be easily digested or metabolized (Table 1). FA-FOS II on simulated digestion by human microbiota released a single metabolite after 48 hours of simulated digestion (Fig 8a). This released metabolite was found to be more polar compared to that of free FA and the UV-VIS spectra were much different from that of FA and FA FOS II. This compound was identified by LC/MS/MS analysis (Fig 8c) as 6-O-p-Coumaroyl-D-glucopyranoside and the m/z of the LC/MS spectra (Fig 8b) matched that of 6-O-p-Coumaroyl-D-glucose. The MS/MS spectra were compared with the spectral data in the NCIM library, and it was found that the spectra matched with that of coumaroyl glucose with an ID score of 0.942 with a mass accuracy of 0.9 (M2 score= 0.97, XIC score = 0.61), the spectral matching and identification was done by using Scaffold Elements, Proteome Software, Inc, Oregon. The parent molecule detected was M-H_2_O-H (i.e 325-19 = 306^-^), in MS /MS the prominent signature fragments detected were 162.02, 145 which matched with that of 6-o-p-Coumaroyl-D-glucose.

**Fig 8.**
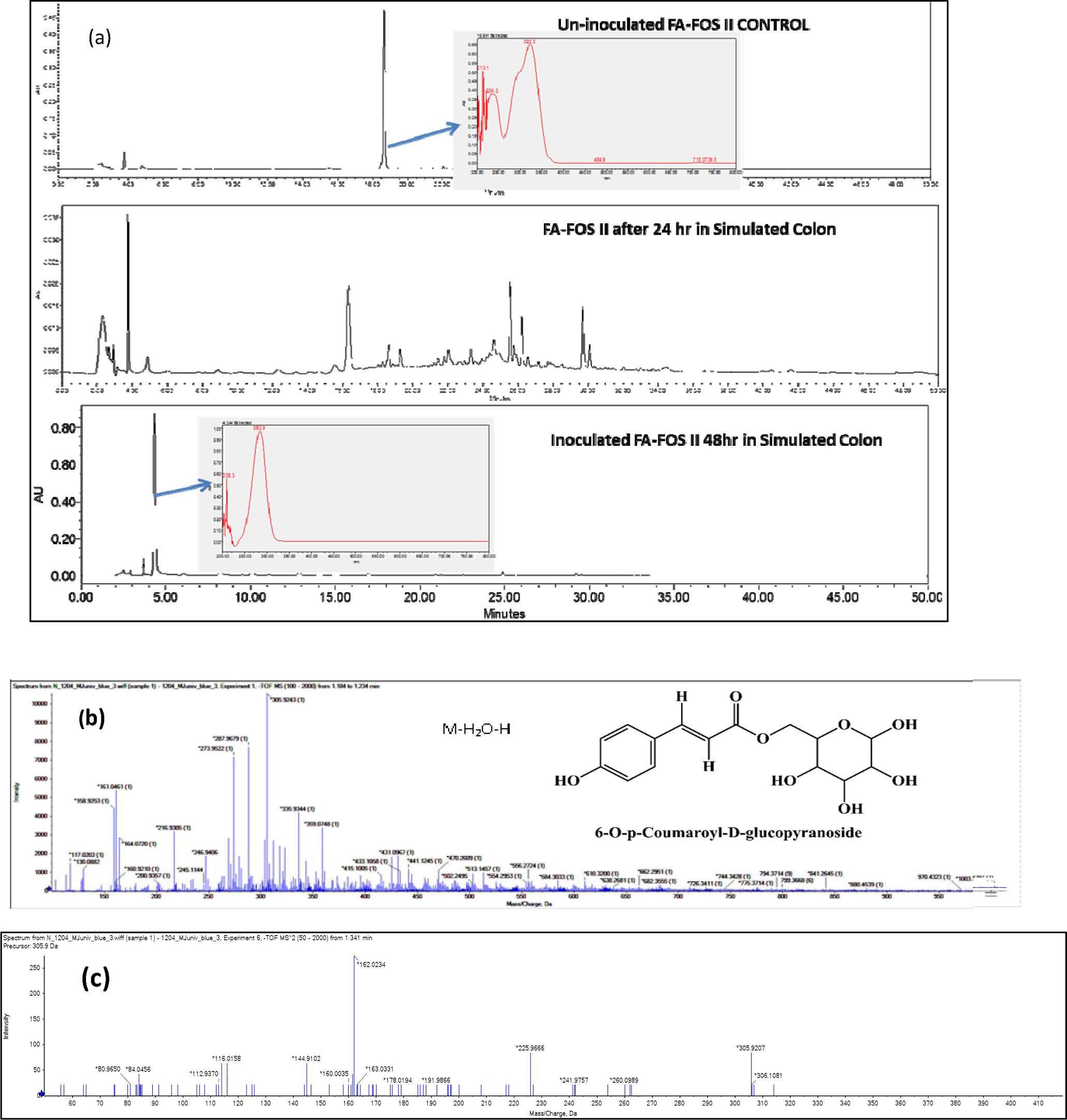
(a) HPLC profile of FA-FOS II after simulated colon digestion at 24 h and 48 h, (b) LC ESI MS of the hydrophilic metabolite, 6-O-p-Coumaroyl-D-glucopyranoside formed from FA-FOS II at 48 h in simulated colon digestion, (c) LC ESI MS/MS of the parent ionic species of m/z 306.

**Table 1:** Metabolic stability and half-life of FA-FOS II in biologically relevant fluids and simulated colonic conditions

| Simulated condition | Stability* (%) | Half-life** (h) |  |
| --- | --- | --- | --- |
| Simulated salivary fluid | 100% | > 4 | 485 |
| Simulated fasted state gastric fluid | 100% | >4 | 486 |
| Simulated fasted state intestinal fluid | 99.96% | >4 | 487 |
| Enteric enzyme cocktail | 54.39% | 36 | 488 |
| Simulated colonic condition | 100% Digested and metabolized | <24 | 489 |

\* Stability was expressed as the release of free ferulate from the conjugate pre- versus post-incubation at 37 °C
\*\* Half-life was expressed as the time at which half of the FA content of FA-FOS I was released into the medium. Data represented as the mean of triplicate experiments

The formation of 6-O-p-Coumaroyl-D-glucopyranoside from FA FOS II was by the catabolic action of the gut microbiota present in the system as inoculum. This catabolic action is known to happen in two steps: one is the demethylation of ferulic acid to caffeic acid and the second is the dehydroxylation of caffeic acid to p-coumaric acid (37). The presence of glucose moiety/ glycoside may be due to the stearic hindrance caused by the electron-dense phenolic acid for the attachment of the feruloyl esterase enzyme which was involved in the catabolic action (37,38). The coumaroyl glycosides were found to inhibit α-glucosidase and α-amylase (39–41), this may be the reason for the cessation of further metabolism of 6-O-p-Coumaroyl-D-glucose to the non-glycosylated end product or other ultimate end product formed in the metabolic fate of such phenolic acids (37,42). The metabolite thus formed from FA FOS II was found to be having anti-oxidant (43) and anti-diabetic properties (40), anti-cancer, anti-inflammatory, anxiolytic, analgesic, and antipyretic activities (44) with mitigatory action against obesity and hyperlipidemia. The biological activities of coumaric acid are found to be enhanced on conjugation with glycosides but their low absorption from dietary sources and hence low bioavailability was the challenge (44). The present mode of delivery and metabolism of FA FOS II holds good prospects as a therapeutic for anti-oxidant, anti-inflammatory, and anti-cancer applications.

### 3.3 Pharmacokinetics of FA FOS II in Rat model

On P.O administration of FA FOS II to rat, the lag time (t_lag_) which is the time delay between the administration of the drug and the detection of the FA above the limits of quantitation in the plasma was significantly high which is 0.25 h compared to that of FA administration via IV and P.O which was 0.083 h. Moreover, the t_max_ for FA FOS II was found to be 0.5 h compared to that of 0.25 h for that an equivalent concentration of FA P.O administration. This along with the t_1/2_ value of FA FOS II shows that the free FA from the FA FOS II is slowly absorbed into the bloodstream after release by the digestive action of the gut microflora (45). The MRT and t_z_ of the free FA from FA FOS II were found to be 4 h and 24 h respectively which is significantly higher when compared to FA IV and P.O administration suggesting that the free FA is slowly and persistently released by the microbial action into the systemic circulation from the large intestine brush border, enabling controlled and sustained release of the payload (FA) at the target site and persistence in the plasma, which increases the exposure of FA in the systemic circulation. However, the plasma bioavailability of FA from FA FOS II was found to be lower which is 20.7% compared to that of FA P.O administration which was found to be 70% which signifies that the majority of FA would be present as conjugated forms (glucuronidated and/or sulphated form) or as their metabolites in the plasma as observed by Azuma et al. (46) and Adam et al. (47). In the postabsorptive phase, the concentration of FA in the plasma was at undetectable limits post administration via I.V and P.O route, the t_z_ of which was 1h and 6h respectively but free FA was found to be detected in plasma up to 24h, this suggests the notion that the enterohepatic cycle does not allow the plasma concentration of FA to remain high in P.O and I.V route of administration. Adam et al. also made similar observations in the in-situ small intestinal perfusion experiment of FA that only 6.2% of the perfused dose of free FA was secreted into bile (47).

Ferulic acid in its free form is found to be quickly absorbed from the stomach and metabolised extensively in the liver and excreted rapidly (48) but a bound form of ferulic acid is absorbed mainly through the large intestine because of the abundance of feruloyl esterase in the lumen produced by the microbiota and are found to be bioavailable for an extended period (49,50). This is due to the reason that free FA could be absorbed by the small intestine very quickly by passive diffusion or by facilitated transport and does not seems to be saturated even at higher luminal concentration (47,51). The glycosylation and conjugation enzymes are abundant in the small intestinal mucosal and submucosal layer compared to that of the large intestine, where in the latter the metabolism and breakdown are modulated by the enzymes secreted by gut microbiota. Hence a large amount of FA absorbed in the small intestine is extensively glucuronidated and found in the serosa of the intestine which is also the common phenomenon with most of the polyphenols (52–55). Thus, the delivery of the FA into the large intestine by FA FOS II has a dual advantage of enhanced exposure of the large intestine lumen with free FA and uptake into the bloodstream without conjugative transformation or extensive glucuronidation.

**Fig. 9.**
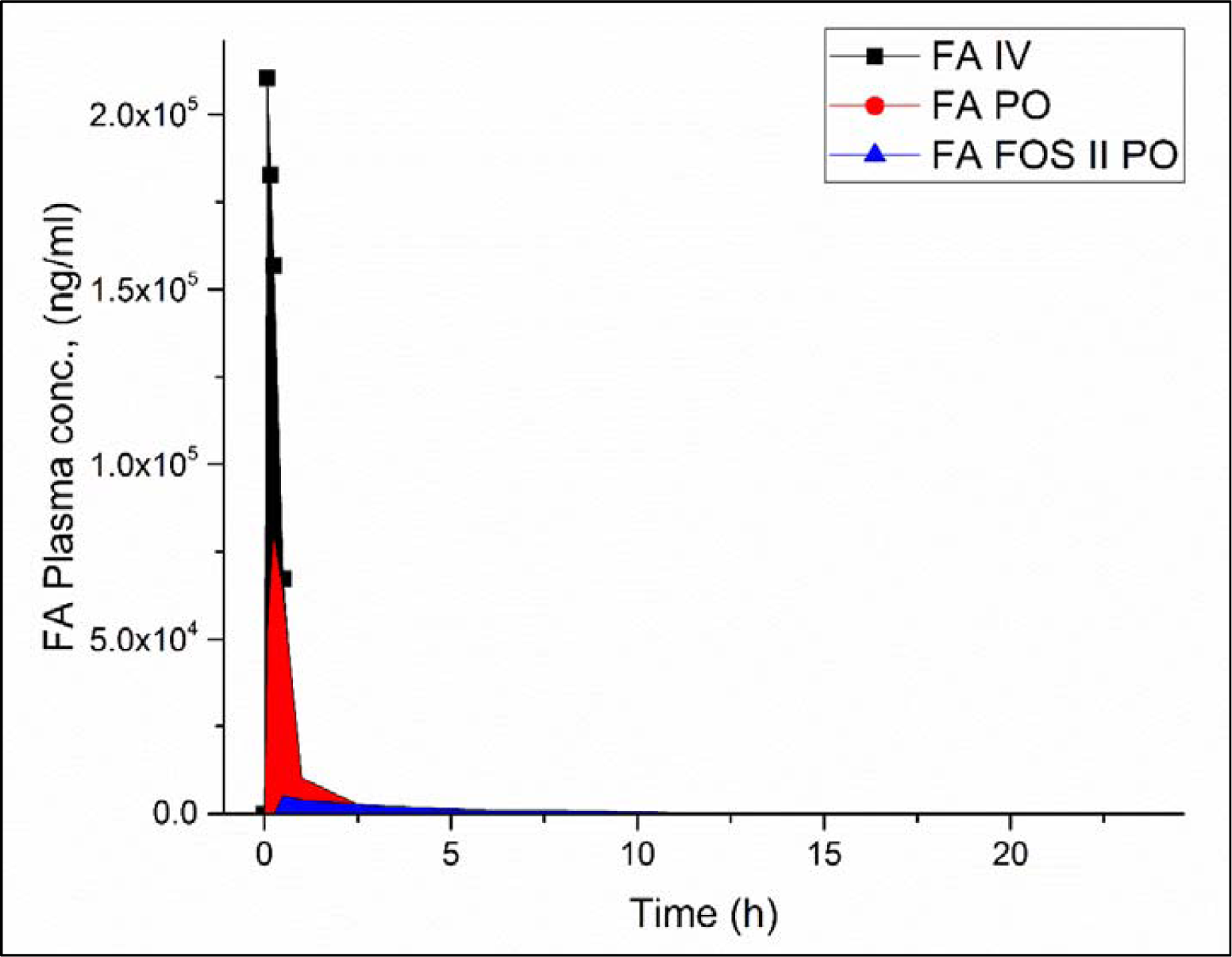
Pharmacokinetic profile of free FA (plasma unbound) in the plasma of rat after IV and oral administration of FA and FA FOS II at 100 mg/Kg body weight free FA equivalent dose.

**Table 2:**
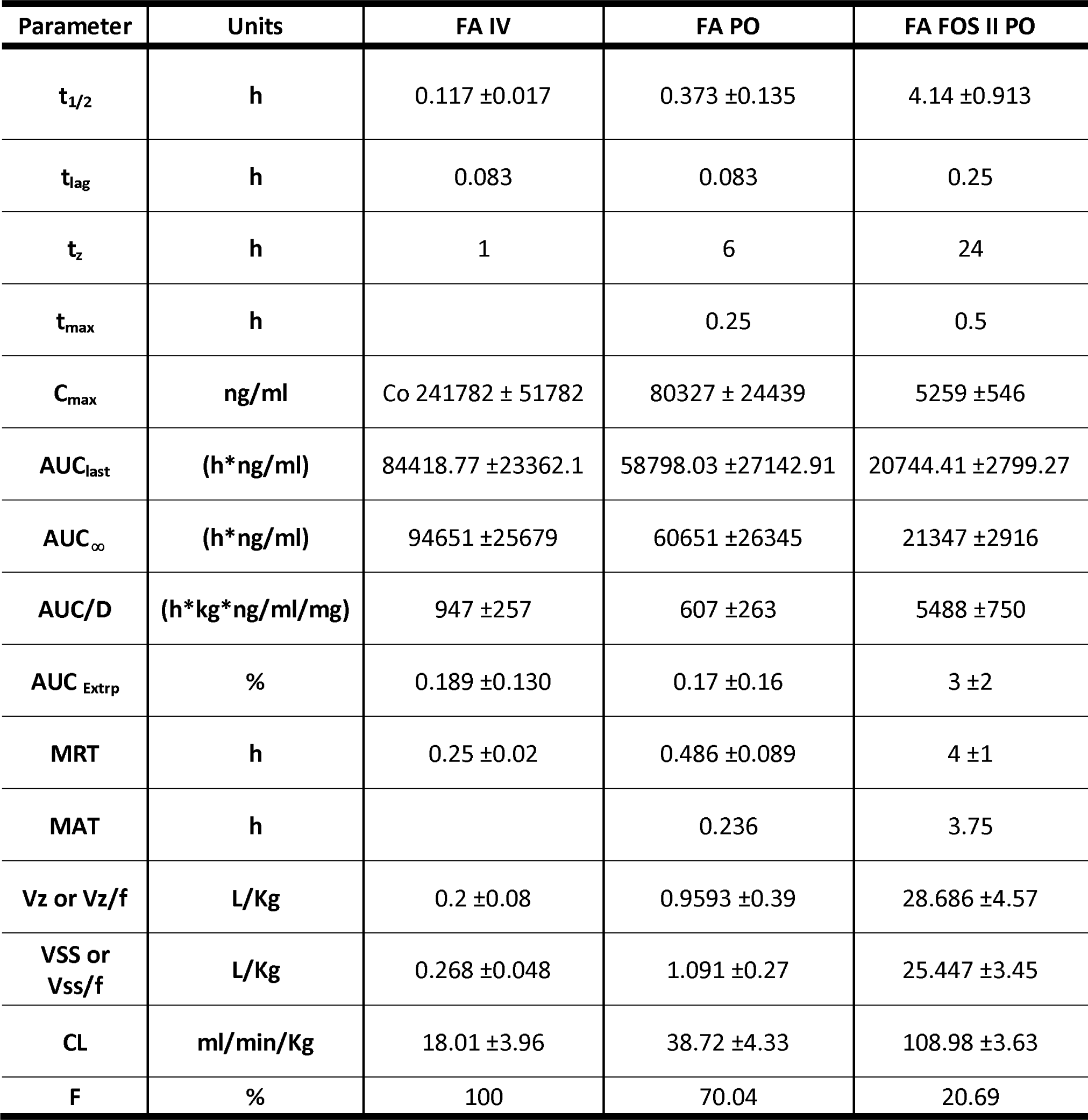
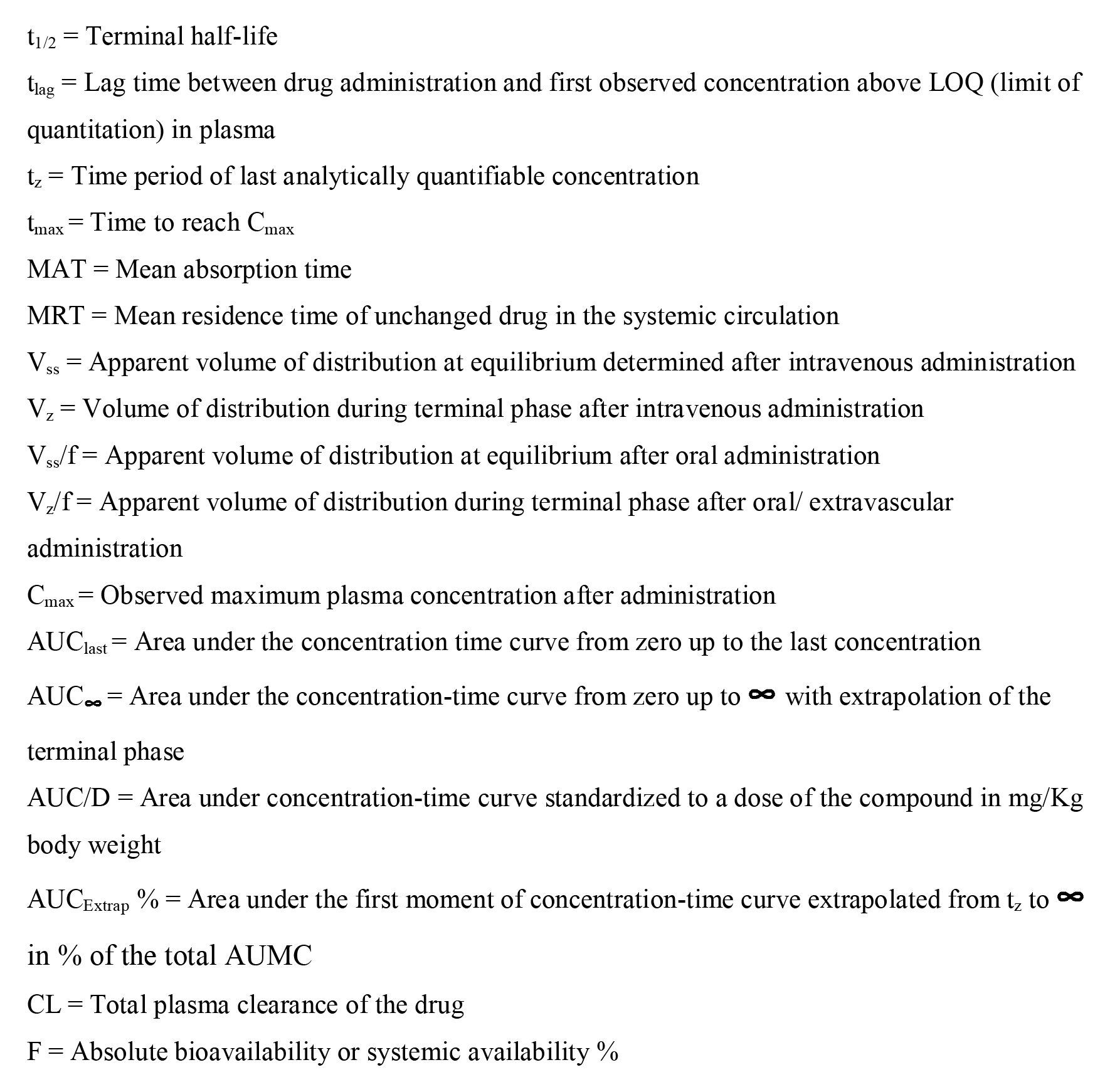
Pharmacokinetic parameters of free FA in the rat plasma

The higher plasma clearance of FA FOS II along with a higher volume of distribution at the equilibrium and terminal phase denotes the higher tissue and cellular distribution of FA in the biological system, this may explain the lower plasma bioavailability. This may be because unlike I.V and P.O administration of FA the administration of FA FOS II generates a higher concentration of non-conjugated non-glucuronidated FA (non-polar molecule or metabolite) in the large intestine or colon which is then uptake into the serosa of the intestine and distributed throughout the periphery of the gut and later into plasma (56–58). It was also reported that more than 49% of FA administrated orally is distributed in the liver and its peripheral tissues (47) and about 53% are found in the gastric mucosa approximately 30min after oral administration and excreted out of these tissues within 60 min of administration (21).

### 3.4 Identification of metabolites of FA FOS II in rat plasma

To determine the various metabolite of FA FOS II apart from the free FA available in the plasma the data-driven parent ion scan was done using a triple quad mass spectrometer. The full MS/MS was performed on the parent and the metabolites were identified as ferulic acid-4-O-sulphate, 1-O-p-coumaroyl-D-glucose, and 6-O-p-coumaroyl-D-glucose, ferulic acid 4-O-glucuronide, 3- (Hydroxyphenyl)propionic acid-O-sulphate, hippuric acid. All these metabolites along with free ferulic acid were detected in the plasma from 30 min to 3h (Table 3), but after this time point, 1-O-p-Coumaroyl-D-glucose and hippuric acid were not detected in the plasma any further. During the terminal phase from 12h to 24h only free ferulic acid and ferulic acid-4-O-sulphate only were detected above LOQ.

**Table 3:** List of metabolites detected at various time points in the rat plasma

| Sampling time | Parent ion/Precursor ion detected |
| --- | --- |
| 0 min | 192.9, 273.1 |
| 30 min | 193.2, 273.2, 325.3, 149.0, 178.0, 229.2, 260.9, 368.9 |
| 1 h | 193.1, 273.3, 325.3, 149.1, 178.1, 229.2, 260.9, 369.3 |
| 3 h | 193.1, 273.2, 325.4, 178.2, 229.1, 261.0, 369.3 |
| 6 h | 193.2, 273.3, 325.0, 369.4 |
| 8 h | 193.1, 273.2, 325.2, 369.3 |
| 12 h | 193.3, 273.0 |
| 24 h | 192.9, 273.0 |

The detected metabolites and their product ions detected were comparable with the previously reported metabolites of FA (59) but sulfoglucuronide of FA was not detected in the plasma after administration of FA FOS II contrary to the previous reports with ferulic acid sugar esters (58). This may be due to the fact that FA FOS II is mainly delivered to the large intestine where the intestinal brush border enzymes are limited in comparison to the small intestine which facilitates the absorption of FA and its metabolites (after microbial action) into the plasma (16) without much glucuronidation by the liver, this may also explain the presence of free FA in the plasma even at 8 to 12 h post administration of FA FOS II.

**Table 4:** The precursor/parent ion detected, and their corresponding product ions observed in the full MS/MS scanning of the parent ion.

| Molecule | Retention time | Monoisotopic mass | [M-H]<sup>-</sup> (m/z) | Product ion m/z detected in our study |
| --- | --- | --- | --- | --- |
| Ferulic acid | 1.83 | 194.057 | 193 | 133.9 |
| Ferulic acid-4'-O-sulphate | 1.83 | 274.014 | 273 | 134, 193 |
| 1-O-p-Coumaroyl-D-glucose<br>(solubility = 8.2 g/L ) | 1.83 | 326.100 | 325 | 134,149 |
| 6-O-p-Coumaroyl-D-glucose<br>(Solubility = 7.67 g/L) | 2.64 | 326.100 | 325 | 266, 279 |
| Ferulic acid 4-O-glucuronide<br>(Solubility = 3.81 g/L) | 1.68, 2.48 | 370.089 | 369 | 193 |
| 3-(Hydroxyphenyl)propionic acid-O-sulphate (Sol = 2.31 g/L) | 2.05 | 262.014 | 261 | 261.00 |
| Hippuric acid (Sol = 1.18 g/L) | 1.81-2.55 | 179.058 | 178 | 133, 178 |

From the XIC chromatogram of the product ions, of various metabolites of FA found in the plasma, it can be assumed that feruloyl sulfate may be the most abundant metabolite in the plasma of rats (Fig.10) since both FA and feruloyl sulfate were detected in the plasma until 24 h post administration of FA FOS II. The order of the metabolite abundance can be represented as Feruloyl-4-O-Sulfate > Ferulic acid > Coumaroyl-D-glucose > Ferulic acid 4-O-glucuronide > Hippuric acid > 3- (Hydroxyphenyl)propionic acid-O-sulphate from their AUC values. This is in line with the previously reported metabolites of FA (20). The present study shows that free FA could be recovered in the plasma and bioavailable even after 12 hours post administration of FA FOS II suggesting a slow and extended-release profile. Glucuronidation and sulfation were the common ways of detoxification or excretion by increasing the polarity/solubility of compounds (20) however the present results are quite different from free FA pharmacokinetics and metabolism. In free FA administration, the FA circulates in the plasma mainly as metabolized form and quickly disappeared from the plasma in 4.5 h after oral administration (20). In the case of 5-O-feruloyl arabinofuranose and feruloyl, arabinoxylan oral administration the total FA concentration (FA and FA metabolites) was detected only up to 4.1 h and the relative plasma bioavailability was found to be 56% and 21% respectively (27). Other reports state that the bound form of ferulic acid such as in wheat bran ferulic acid was detected in the plasma up to 24 h (56). This supports the notion that the targeted delivery to the large intestine could facilitate the extended-release of FA in a controlled manner without much metabolic transformation. This difference in bioavailability and pharmacokinetic profile may be due to the difference in the form of FA (bound to polysaccharide or sugar esters) and the oral dose which affects the ultimate fate of FA in the gastrointestinal tract and its systemic bioavailability (60). Absorbability is an important factor that describes the fraction of administered dose in the formulation or agent that traverses the intestinal mucosal barrier which can also partly determine the bioavailability of FA. It is suggested that the order of absorbability of dietary FA is as follows: free FA > Feruloylated sugars/mono-saccharides > Feruloylated polysaccharide (16,58). This may be because the complex structure of the oligosaccharide and polysaccharides in the feruloylated structure reduces the interaction or binding of hydrolyzing enzymes such as feruloyl esterases to such polymeric or complex conjugates of ferulic acid and thus reducing the rate of release of FA (57). Thus the form of feruloylated molecules plays a vital role in the absorbability and subsequent bioavailability and excretion of FA in rats and humans (47,57,60).

**Fig. 10.**
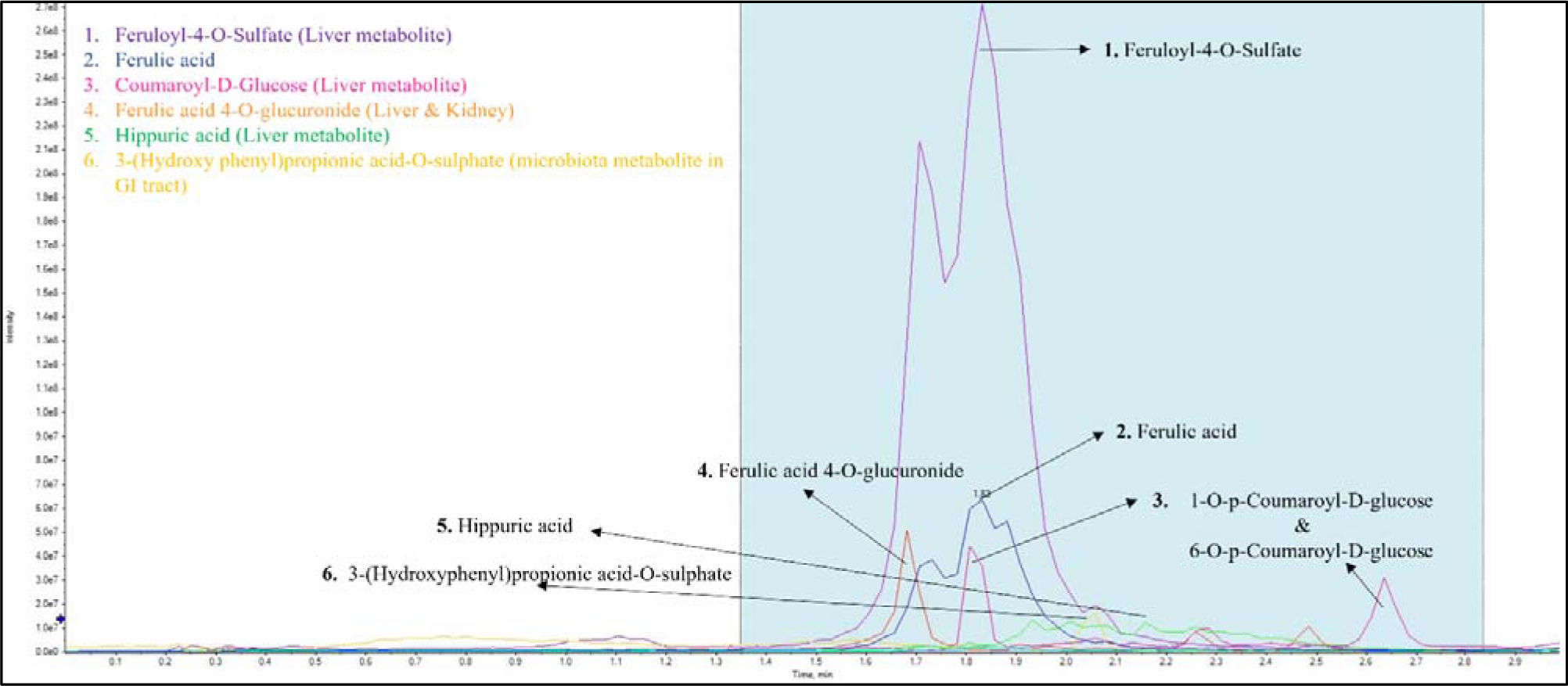
Representative XIC chromatogram of the production of various metabolites found in the plasma of rat at 30 min after administration of FA FOS II

### 3.5 Efficacy studies in Xenograft tumour carrying mouse model

To determine the bioavailability and efficacy of FA by its distribution in the tumour tissue on oral administration of FA FOS II, a xenograft mouse model carrying tumour from human colorectal cancer cell line HT-29 with P.O administration of 100 mg/Kg body weight per day FA equivalent of FA FOS II, for a period of four weeks was carried out and investigated for the amelioration of tumour development. From the xenograft model efficacy study, it was observed that there was a considerable 50% reduction in the tumour volume in the FA FOS II treatment group compared to the control group (p<0.05), this was comparable to 52.50 % reduction in tumour volume by 5-FU group despite their adverse side effects. This result was reflected well in the tumour weight data at the termination of four weeks of treatment, in which FA FOS II showed a 37.34% reduction in the mean tumour weight compared to the control group. The tumour volume growth rate profile shows a significant decrease in the tumour proliferation of 56.8% per day (p<0.05) when compared to the control group. When we gauge the toxicity or adverse side effects of FA FOS II treatment, they show no drop in the body weight of the mice but rather a 6% increased weight gain when compared to the control group. There was a 41.47 % increase in the daily body weight gain rate in FA FOS II compared to the control group. In the body weight gain profile during the treatment regime, FA FOS II scored higher than the control group during the entire treatment period suggesting no adverse reactions to FA FOS II, while in the case of 5-FU body weight steeply decreased during the initial treatment regime, and a steady rise in the body weight during the resting phase indicating the adverse effect of 5-FU on the mice physiology.

**Fig. 11.**
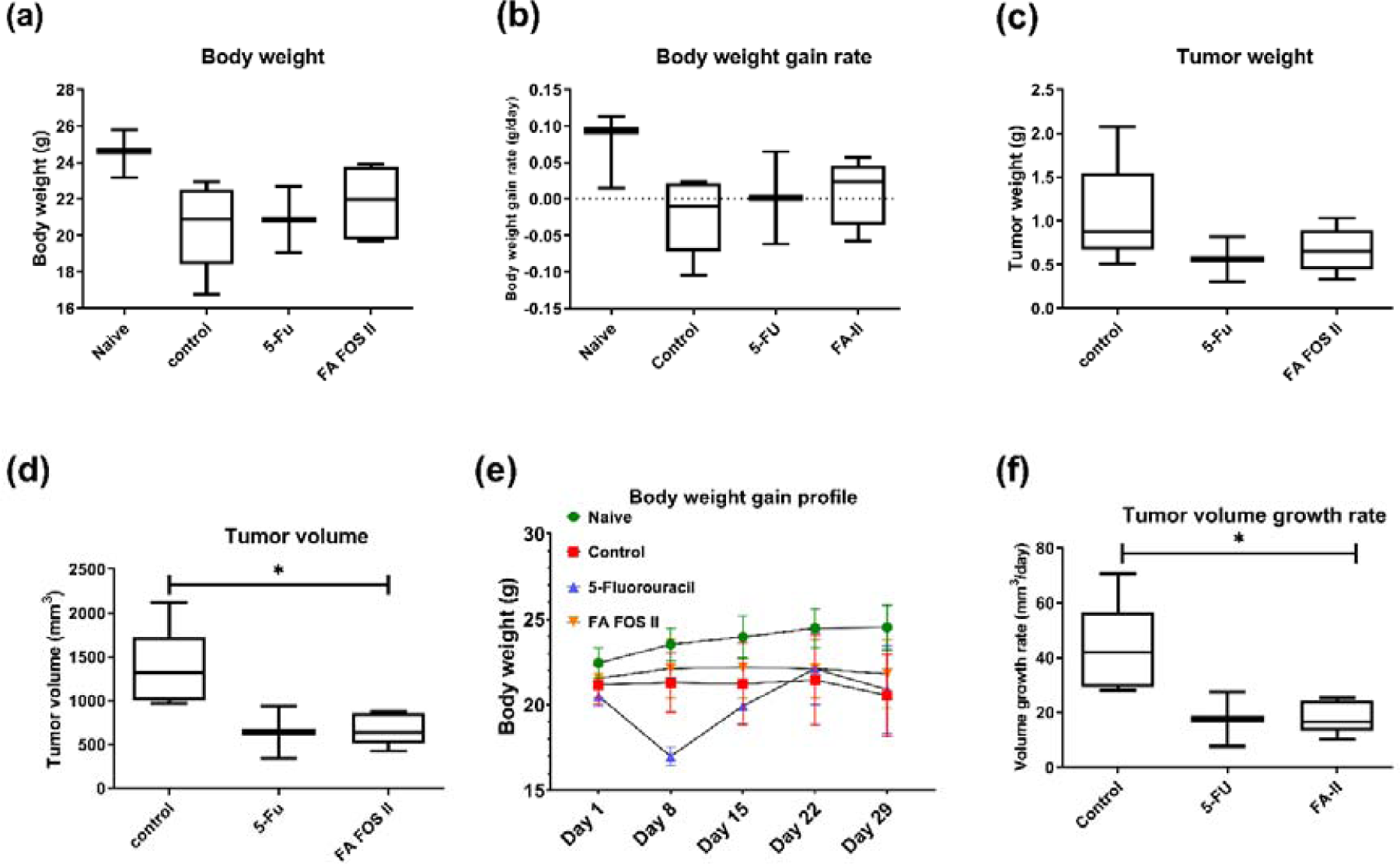
Macroscopic parameters in Xenograft mice carrying tumors from HT-29 human colon cancer cells treated with FA FOS II and 5-fluorouracil for a period of four weeks. (a) Body weight after four weeks of treatment, (b) Body weight growth rate (g/day), (c) Tumour weight (g) after the elapse of a four-week treatment period, (d) Tumour volume after four weeks of the treatment period, (e) Body weight gain profile over the period of four weeks treatment period, (f) Tumour volume growth rate (mm^3^/day) over the period of four weeks.

The histopathological examination of the core portion (longitudinal) of the excised tumour hematoxylin and eosin staining revealed that the tumour occupied regions in the FA FOS II treatment group were 32.77 %/mm^2^ (±1.03) compared to that of the Control group which had a tumour occupied region of 49.32 %/mm^2^ (± 5.73). The 5-FU group had a tumour occupied region of 33.50 %/mm^2^ (± 0.56) comparable with that of FA FOS II. The scanned slides were then analyzed by using Qpath 0.3.2 digital pathology and image analysis software to determine the tumour cells, necrotic cells, and immune cells in the excised tumour mass (Fig. 13).

**Fig. 12.**
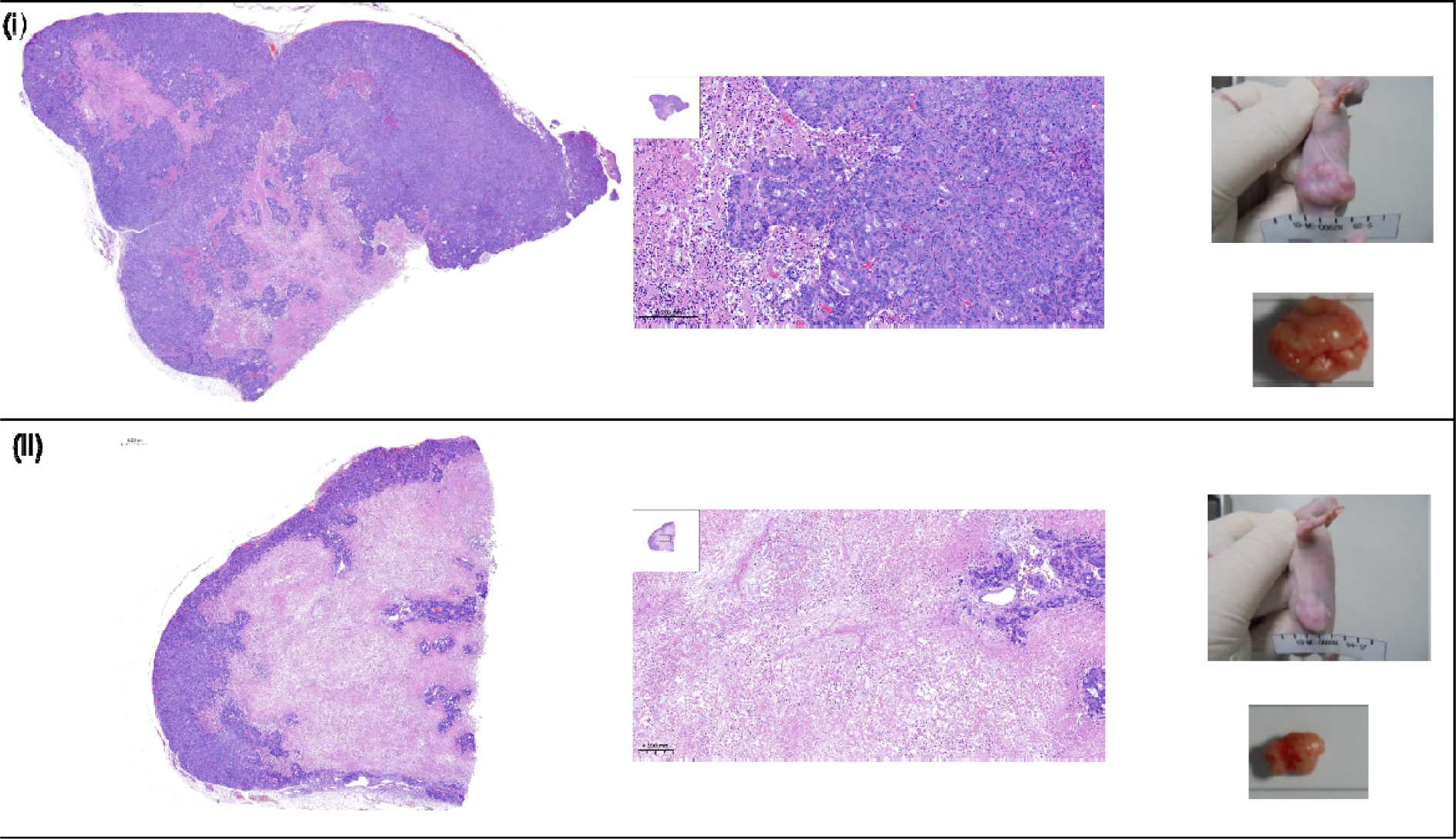
Hematoxylin and Eosin staining of excised tumour from control mice and FA FOS II treated mice, inset-showing 40x objective, far-right images-Representative picture of mice bearing xenograft tumour and the excised tumour from each experimental group, control and FA FOS II treatment group after four weeks treatment respectively.

**Fig. 13.**
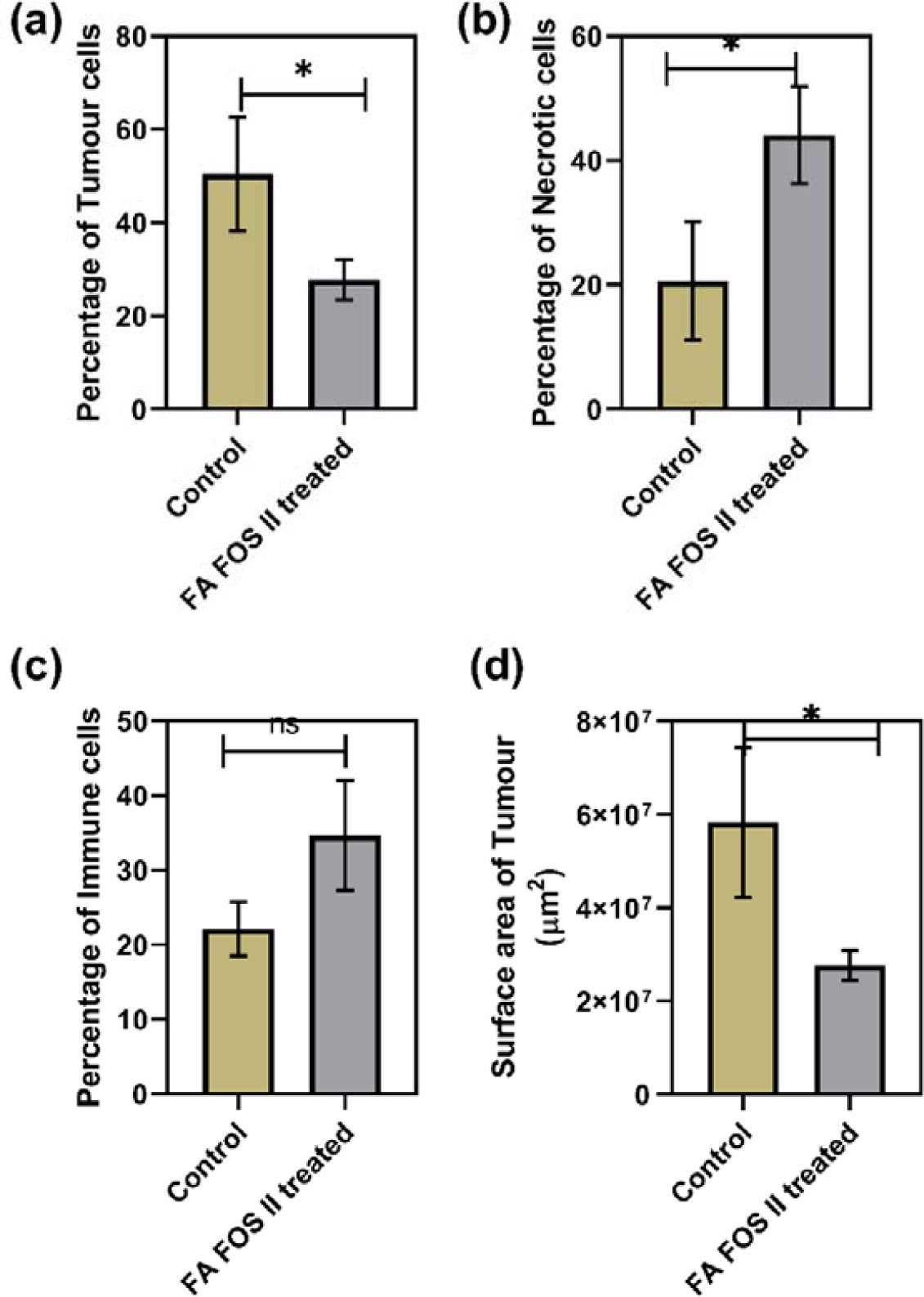
Histological score of the H&E-stained excised xenograft tumour tissue determined after 4 weeks of treatment with FA FOS II. (a) Percentage of tumour cells occupying the xenograft tumour, (b) Percentage of necrotic cells, (c) Percentage of immune cells (macrophages, neutrophils, basophils, monocytes, eosinophils) in the tumour occupied region

The digital pathology using Qpath revealed that there was a 44.98 % reduction (p<0.05) in the tumour cells in the H&E slides in the FA FOS II treatment group compared to the control group. In the control group, 50.44% ±12.21 cells were tumour cells compared to 27.75% ±4.32 tumour cells in the FA FOS II group. There was a 114% (p< 0.05) increase in the necrotic cell population in the FA FOS II treatment group compared to the control group, this also concurs with the minimal chromatic void space-like region found in the H&E-stained FA FOS II histopathology slides of excised tumors (Fig. 12). There was a 56.2% increase in the immune cells found surrounding the tumour micro-environment in the FA FOS II treatment group compared to the control group. On average there was a 52.5% reduction in the surface area of the (central core) microtome section of the excised tumour tissue of the FA FOS II treatment group compared to that of the control group. To determine the expression of various proteins responsible for the reduced proliferation and cell death in tumour cells, immunohistochemistry (IHC) was performed and the cellular localization of the target proteins with relevant antibodies was determined and analysed. To confirm the apoptosis/necrosis TUNNEL assay (Immuno Fluorescence) was performed to validate the results.

**Fig. 14.**
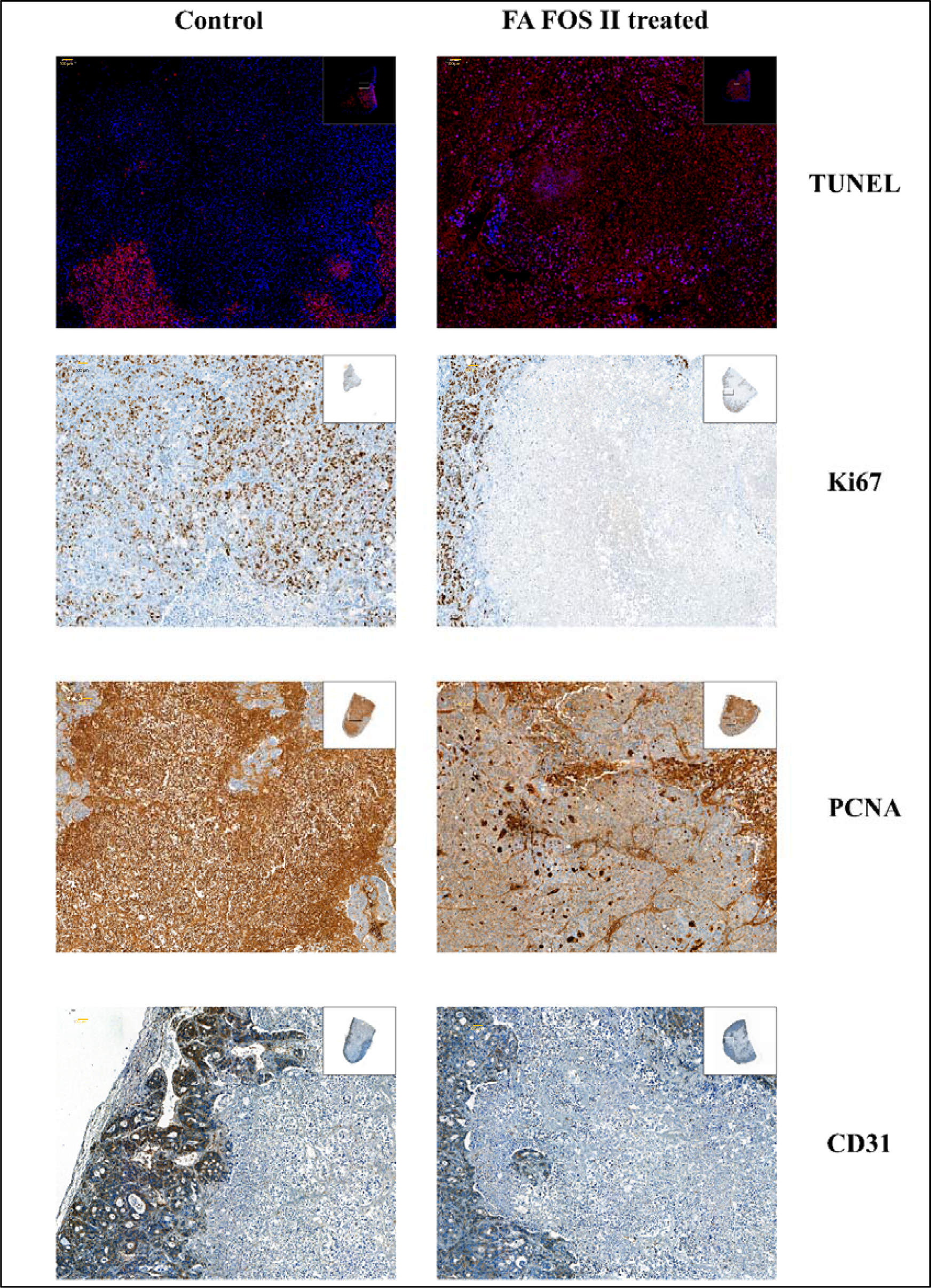
Photomicrograph showing TUNEL Immunofluorescence, Nuclear localization, and expression of Ki67, p53, and PCNA, cytoplasmic expression of CD31 mouse antigen (angiogenesis-related protein) Immunohistochemistry (Hematoxylin and DAB).

**Fig. 15.**
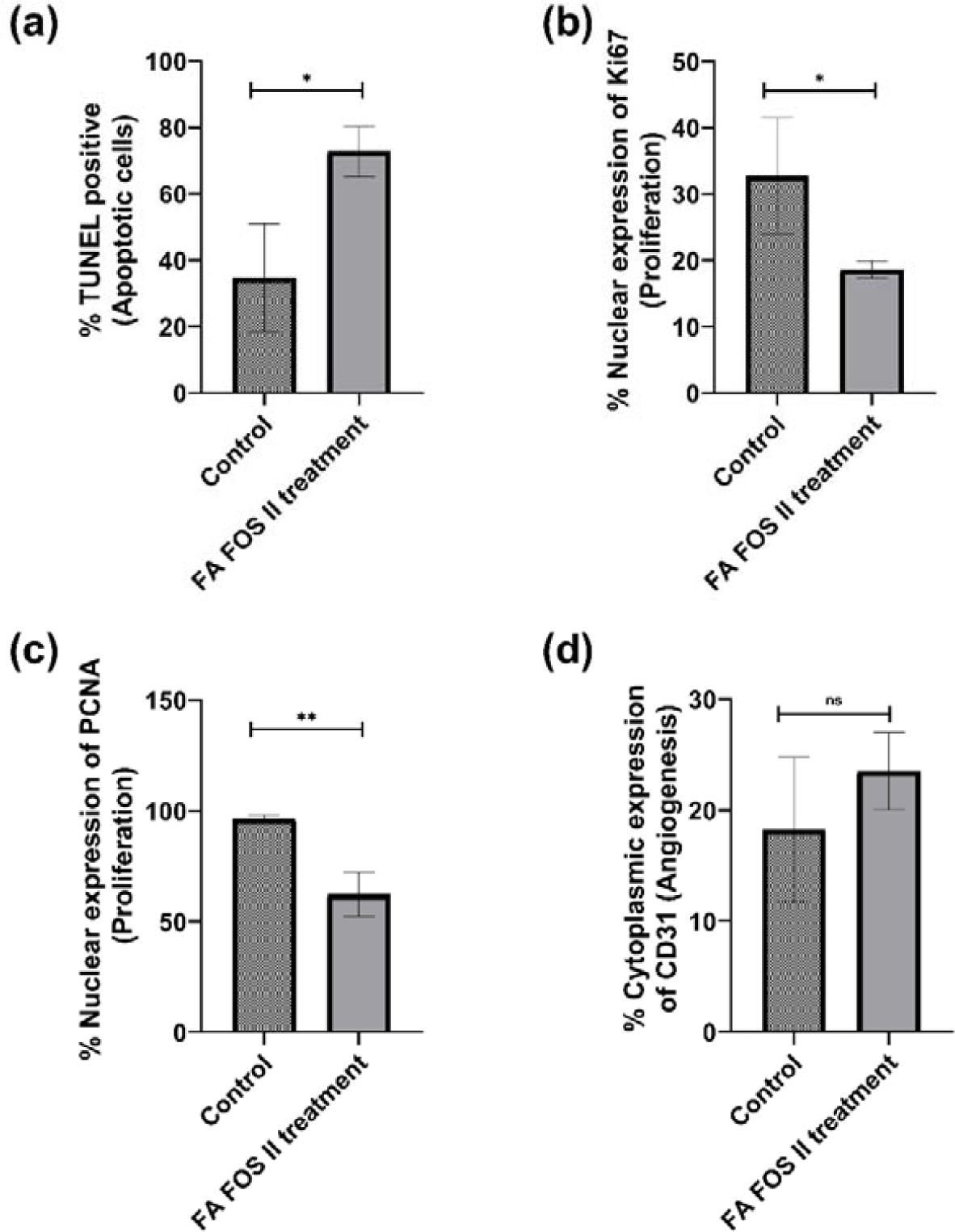
Scoring of (a) TUNEL positive cells, (b) Nuclear Ki67 expressing cells, (c) Nuclear PCNA expressing cells, (d) Cytosolic expression of CD31

DNA fragmentation is the hallmark of apoptosis, the terminal deoxynucleotidyl transferase dUTP nick end labelling (TUNEL) is used to detect and measure the cells which underwent DNA fragmentation and hence measuring the extent and apoptotic phase determination. In this study the nuclei were labelled with DAPI and BrdU labelled with red fluorophore by using a TUNEL assay kit of abcam. The total number of cells in the tumour was determined by counting the nuclei of cells with DAPI fluorescence and the cells undergoing apoptosis and DNA fragmentation were determined by counting the cells with BrdU red fluorescence in the nucleus. It was found that there was a 110% increase in the TUNEL-positive cells in the tumours treated with FA FOS II compared to that of the control group. The FA FOS II treatment group showed 72.73% ± 7.6 TUNEL positive cells while the control group had only 34.6% ±16.2 TUNEL positive cells. The nuclear expression of the Ki67 antigen is a reliable marker of tumour cell proliferation (61)and serves as an prognosis tool for colorectal cancer (62). Ki67 was present only during the active phase of the cell cycle and was absent in the resting cells and the expression of Ki67 protein is associated only with proliferating cells of malignant tumours since the half-life of this protein is only 1-1.5 h in normal cells the continual expression and nuclear localization are found to be associated with tumour aggressiveness (62). In this study, the nuclear localization and expression level of Ki67 was determined in each experimental group and quantified. It was found that there was a 43.12% reduction (p<0.05) in the nuclear Ki67 expression in the FA FOS II treatment group compared to that of the control group. The control group had 32.7% ±8.8 of cells expressing Ki67 in the nucleus. Proliferating cell nuclear antigen (PCNA) was known to be a cyclin or ancillary protein for DNA polymerase delta and it was found to be localized into the nucleus. The expression level was found to be elevated during the replication process of DNA and plays a major role in the proliferation of cells. Its expression along with the co-expression of other markers plays a prime role in cell division and proliferation, and hence their elevated expression was considered a predictor of poor prognosis in colorectal cancer (63). In this study with a xenograft mouse model carrying a tumour from a human colorectal cancer cell line, there was observed a 35.35% reduction (p<0.005) in the nuclear localization and expression of PCNA in the FA FOS II treated group compared to the control group. The control group showed 96.33% ± 1.74 of cells positive for nuclear localization while the FA FOS II treatment group showed 62.27% ±9.8 of cells positive for PCNA nuclear expression. CD31 are considered to be among the prognostic angiogenic marker which is considered to be the main mediator of tumour angiogenesis. Tumour angiogenesis is considered to be a poor prognosis in patients with gastrointestinal stromal tumours (GIST) which are the prevalent mesenchymal tumours of the gastrointestinal tract (64–66). Angiogenesis is considered a contributor to the growth and spread of gastrointestinal tumour. Here in this study, the expression level of mouse CD31 in the xenograft tumour mass is determined to evaluate whether there is any pro-angiogenesis or suppression of angiogenesis activity happening in the blood vessels feeding the tumour. It was found that there was no statistically significant difference found in the angiogenic marker CD31 expression level between the control group and FA FOS II treatment group.

### 3.6 MALDI imaging: determination of active metabolite of FA FOS II and validation of targeted delivery

MALDI imaging of the tissue is employed in determining the spatial distribution, identification, and quantitation of proteins, small molecules, lipids, and metabolites in the tissue samples. They are capable of generating a lot of biologically relevant data useful in determining the exposure level, metabolism, tissue uptake, and delivery of active drugs or molecules (67).

From the MALDI imaging and differential distribution analysis in the xenograft mouse model carrying human HT-29 colorectal tumour. It was evident that FA is made bioavailable to the tumour tissue and the accumulation of the same in the form of the most abundant metabolite ferulic acid sulphate, within the tumour tissue, concurring with the plasma metabolite data of rats (Fig. 10). It was also found that ferulic acid sulphate (FAS) was preferentially uptake and distributed into the tumour tissue (Fig.16). The FAS which is the metabolite of FA formed in the liver due to the conjugation action of liver enzymes rendered the molecule water-soluble and thereby aiding in the transport and distribution into the tissues away from the absorption site which is the intestine (68). It is also observed that feruloylated sulfate and glucuronide demonstrated larger physiological effects than free FA (69,70). The sulfation of FA takes place mainly in the liver through the activity of sulfotransferase. It was reported that FAS was found to be in the circulatory system and a major metabolite of FA was found in the celiac artery (17). Orally administered conjugated FA is known to provide a lasting and consistent presence of FA due to its gradual absorption in the gastrointestinal tract(17). In xenograft tumour tissue ferulic acid sulphate was detected as a sodium and potassium adduct (Table 5). In the AOM DSS mediated colitis induced colon cancer mouse model FA accumulated specifically into the aberrant and tumour tissue as ferulic acid sulphate. The FA and its metabolites are known to be transported into the intestinal mucosa towards the serosa up to 90% due to passive diffusion and also aided by monocarboxylic acid transporter and carrier-mediated transport mechanism (17). The FA treatment group showed no presence of FA or its metabolites suggesting poor bioavailability of FA in the larger intestine when administered orally. This may be because most of the free FA is absorbed rapidly in the stomach and extensively metabolized by the liver and excreted quickly from the system (17). The liver tissue of the FA FOS II treatment group showed the presence of ferulic acid glucose (ferulic acid glucuronide) and the tissue distribution was comparably abundant in the FA FOS II treatment group compared to the FA treatment group (Fig.17 (b)). Ferulic acid glucuronides are usually liver metabolite but in the FA FOS II the action of the gut microbiota was found to release ferulic acid glucose which in turn is found to be de-carboxylated to coumaroyl glucose in the large intestine (Fig. 8). Since they are already in the glucuronidated form the absorbed ferulic acid glucuronide may be transported into the bloodstream without any further metabolism by the liver. The higher presence of ferulic aid glucose in the liver denotes sustained and consistent release of ferulic acid glucose in the large intestine and uptake into the bloodstream. Previous reports suggest that more than 49% of intestinal perfusion of free FA is distributed in the liver, kidney, and their peripheral tissues (47) but in this study of administration of FA FOS II the metabolite of FA and free FA released from the oligosaccharide in the intestine are found to be present in the plasma, subcutaneous tissues and distributed in the intestinal tissues for an extended period compared to free FA administration. This may be because the glucuronidated FA released and absorbed in the lower gastrointestinal tract could evade further first-pass metabolism by the liver and be made available in circulation.

**Fig. 16.**
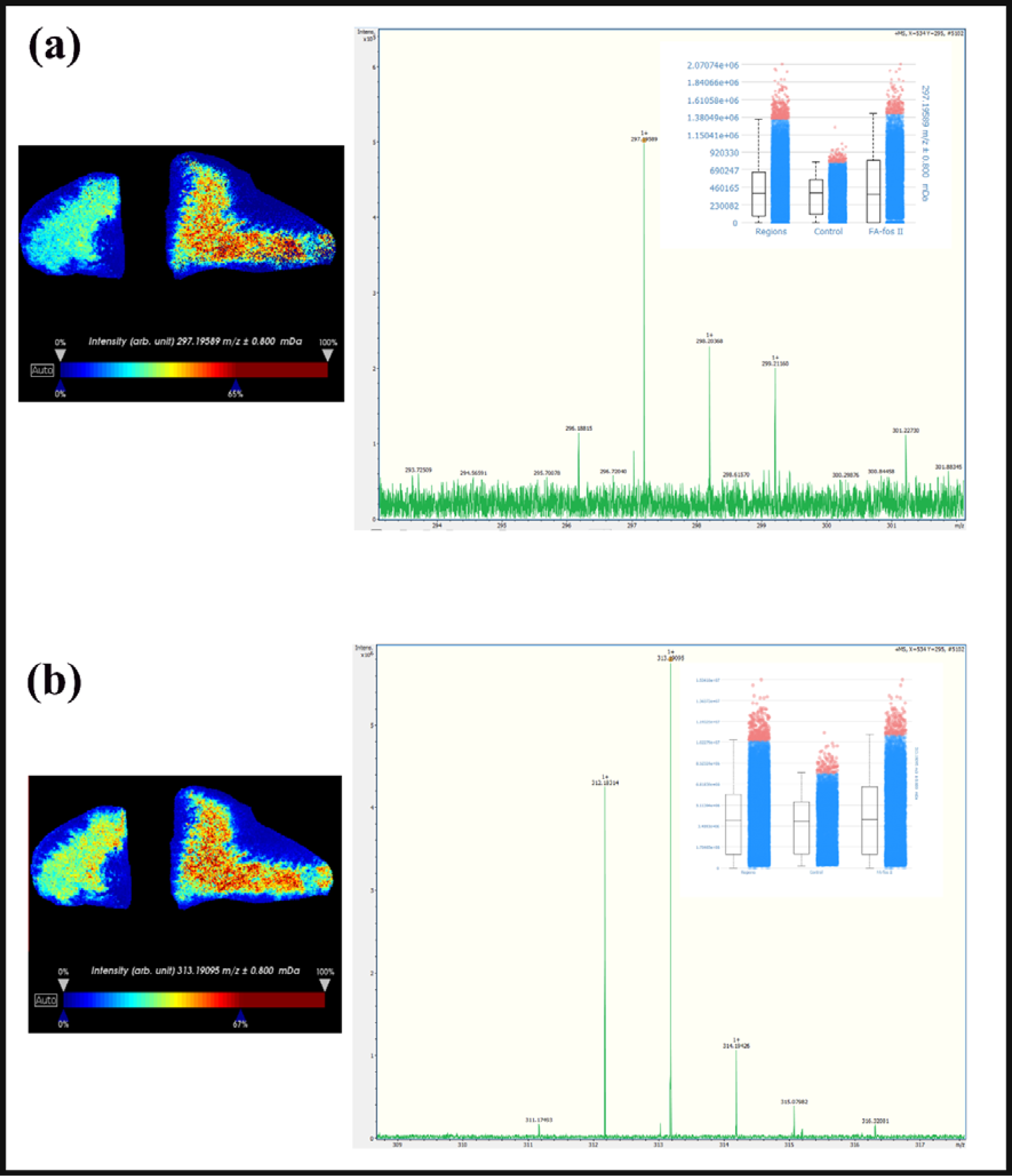
MALDI imaging of xenograft tumour from nude mice after 4 weeks of treatment (6 h post-FA FOS II administration). The left image depicts the metabolite abundance and distribution, the control group tumour is on the left and the FA FOS II treated xenograft tumour is on the right. The far-right side image depicts the MS spectra of the metabolite, inset-the differential abundance of the metabolite, MS score between the control group and FA FOS II treated tumour.

**Fig. 17.**
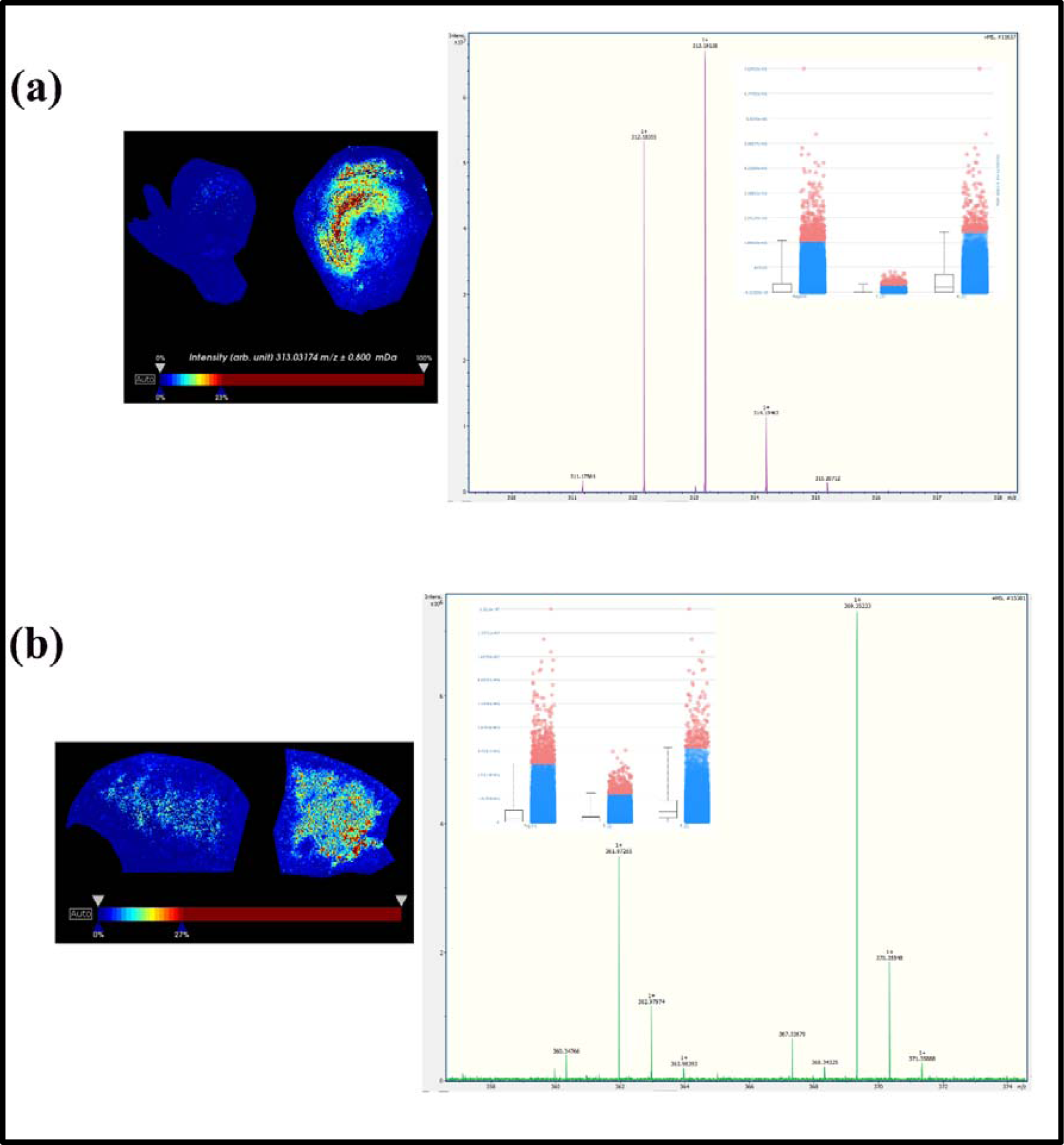
MALDI imaging of (a) Full-length colon (swiss roll) and (b) liver section of AOM DSS mediated colitis induced colon cancer mouse model after 4 weeks treatment (6 h post-FA and FA FOS II administration). The left image depicts the metabolite abundance and distribution, the FA-treated tissue/organ sample is on the left, and the FA FOS II treated tissue/organ sample is on the right of the photo plate. The far-right side image depicts the MS spectra of the metabolite, inset-the differential abundance of the metabolite, and MS score between the FA treatment and FA FOS II treatment group.

**Fig. 18.**
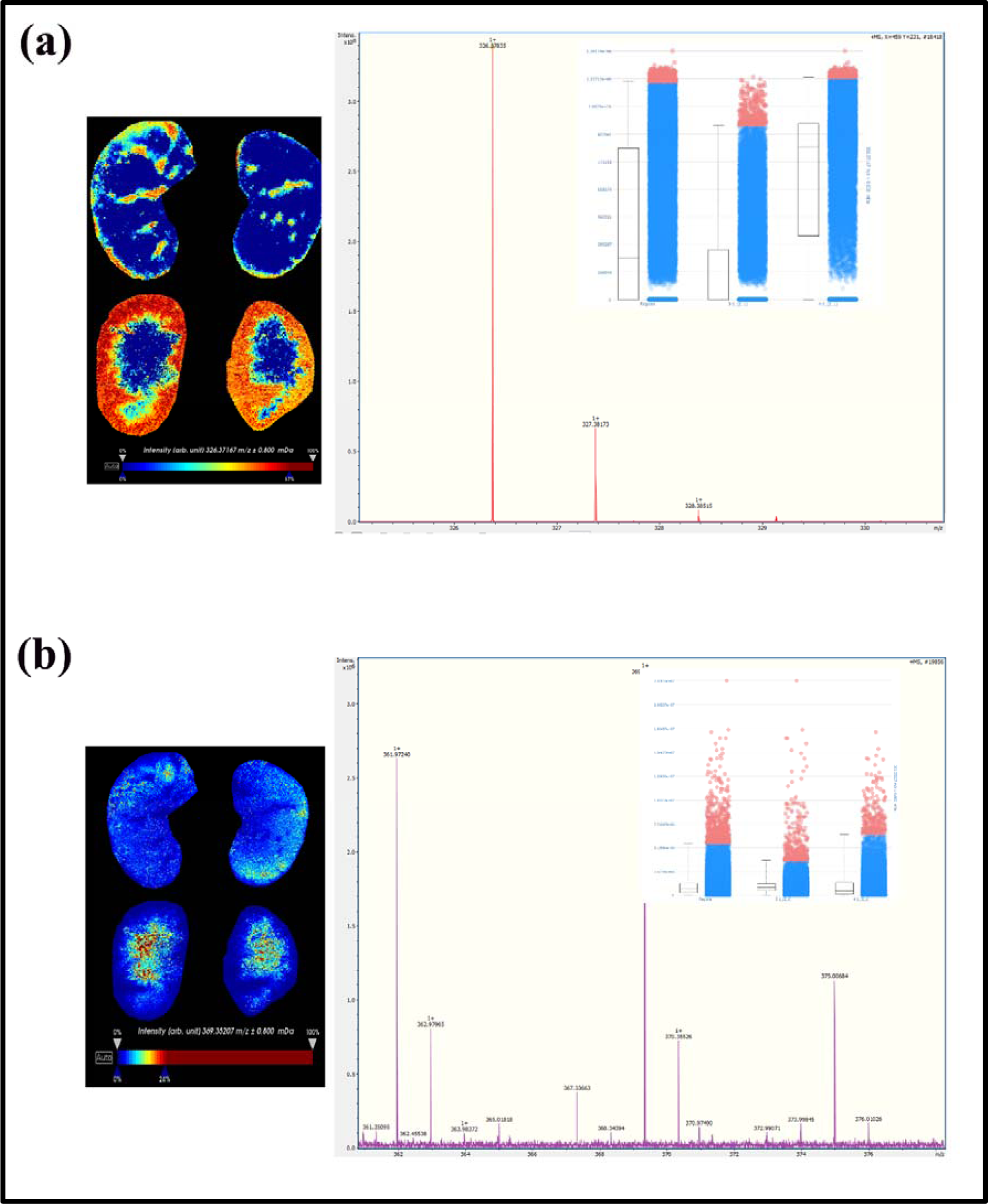
MALDI imaging of Kidney sections of AOM DSS mediated colitis induced colon cancer, mouse model. The left image depicts the metabolite abundance and distribution, the FA-treated tissue/organ sample is on the top and the FA FOS II treated tissue/organ sample is on the bottom of the photo plate. The far-right side image depicts the MS spectra of the metabolite, inset-the differential abundance of the metabolite, and MS score between the FA treatment and FA FOS II treatment group.

**Table 5:** The metabolites of FA FOS II detected in the tissue/organ six hours post administration of FA FOS II, tissue localization, and differential distribution

| Tissue | Metabolite<br>(Monoisotopic<br>mass) | Adduct | Detected m/z | AUC<br>(False positive<br>vs<br>True positive) |
| --- | --- | --- | --- | --- |
| Xenograft tumour | Ferulic acid 4'-O-Sulphate<br>(274.01) | Na adducts<br>(296.99) | 297.19 ±0.8 mDa | 0.527 |
| Xenograft tumour | Ferulic acid 4'-O-Sulphate<br>(274.01) | K adducts<br>(313.1) | 313.19 ±0.8 mDa | 0.555 |
| AOM DSS colon | Ferulic acid 4'-O-Sulphate<br>(274.01) | K adducts<br>(313.1) | 313.19 ±0.8 mDa | 0.844 |
| AOM DSS Kidney | Coumaroyl-D-glucose<br>(326.10) | - | 326.37 ±0.8 mDa | 0.820 |
| AOM DSS Kidney | Ferulic acid<br>glucuronide<br>(370.08) | - | 370.35 ±0.8 mDa | 0.405 |
| AOM DSS Liver | Ferulic acid<br>glucuronide<br>(370.08) | - | 370.35 ±0.8 mDa | 0.826 |

The kidney tissue sections depicted the presence of ferulic acid glucose and coumaroyl glucose which is the end product or final metabolite form available in the plasma to be excreted in the urine. Both are metabolites of microbiota activity in the intestine. Among the two metabolites detected in the kidney coumaroyl glucose was found to be the most abundant end product in the mouse kidney. This may suggest that coumaroyl glucose could be the largest metabolite in the system after ferulic acid sulphate and free ferulic acid which also concurs well with the rat plasma metabolite data (Fig. 10). Coumaroyl glucose could be one of the active metabolites of FA FOS II available in the plasma and in other tissues made available through the systemic circulation. Previous reports also suggest that ferulic acid is mainly excreted through urine mainly as conjugated forms (16,48,51). Reports also suggest that in the urinary excretion of FA from the conjugated forms such as in wheat bran the elimination rate was found to be slowed by 15 fold than the consumption of free FA (19,20)

## 4. Conclusion

The synthesized FA FOS II was found to be water-soluble, resistant to gastric digestion and absorption, amicable to digestion by intestinal microbiota, and release free ferulic acid in the large intestine. They are found to release free ferulic acid consistently for an extended period and thereby increasing the exposure of FA and its metabolites in the plasma, large intestine, and in tissues away from the site of delivery. They are found to be accumulated specifically in considerable amounts in the tumour tissue far from the intestine (delivery site) and also localized in the tumour cells and cancerous tissues in the colon. Thus, this method of oral drug delivery systems could aid in the development of targeted delivery to the large intestine or colon where the water activity is low. These systems could also reduce the dosage and interval of drug administration needed to maintain a specific level of drug in the plasma since they are found to be consistent in the release of the payload/drug for an extended period. These conjugated systems could also be used as a tool to bypass liver metabolism and/or metabolism by upper intestinal enzymes enabling the delivery of the drugs in the intended form without much loss to metabolism and rapid excretion.

## Funding

This work is carried out with the support of the “Cooperative Research Program for Agriculture Science & Technology Development (Project No. PJ015624 assigned to Joo-Won Suh)”, the Rural Development Administration of the Republic of Korea.

Eldin M Johnson thankfully acknowledges the fellowship grant from the Department of Biotechnology (BT/PR6486/GBD/27/433/2012), Govt. of India, New Delhi, India.

&

“Cooperative Research Program for Agriculture Science & Technology Development (Project No. PJ01319101, PJ011287022016)”, the Rural Development Administration of the Republic of Korea.

## Author contribution

**Eldin M. Johnson:** Conceptualization, Design of study, Methodology, Investigation, Formal analysis, Data curation, Software, Writing - original draft and editing. **Joo-Won Suh:** Supervision, Resources, Project administration, Funding acquisition, Editing.

